# A new mRNA antigen vaccine induces potent B and T cell responses and *in vivo* protection against SARS-CoV-2

**DOI:** 10.64898/2026.03.02.709177

**Authors:** Jing Wen, Jaesu Moon, Luca Tucciarone, Te-Hsuan Bu, Amanda Y. Sun, Robyn Miller, Julia Timis, Lujing Wu, Davey M. Smith, Sujan Shresta, Kyle J. Gaulton, Tariq M. Rana

**Affiliations:** Department of Cellular and Molecular Medicine, University of California San Diego, 9500 Gilman Drive, La Jolla, California 92093, USA; Department of Pediatrics, University of California San Diego, 9500 Gilman Drive, La Jolla, CA 92093, USA; Center for Vaccine Innovation, La Jolla Institute for Immunology, 9420 Athena Cir, La Jolla, CA, 92037 USA; Department of Medicine, University of California San Diego, La Jolla, California, USA

## Abstract

The SARS-CoV-2 mRNA vaccine provides effective protection against viral infection and severe disease by inducing efficient adaptive immunity. However, vaccine efficacy is decreased against emerging variants, and immune memory is relatively short-lived. Here, we added new T cell epitopes to the RBD (receptor-binding domain) mRNA vaccine and identified a SARS-CoV-2 membrane epitope that significantly improved vaccine-induced immunity and protection in vivo. That new vaccine, designated G1-C, induced 8.2-fold higher levels of RBD-specific antibodies than did RBD and enhanced spike-specific T cell and B cell responses. Remarkably, the G1-C modulated hematopoietic stem cell (HSC) differentiation and increased levels of B and NK cells by regulating multiple signaling pathways in bone marrow potentially via Fos, Klf4, and Klf6 transcription factors. Altogether, these findings identify a new vaccine candidate to control viral infection by affecting the lymphoid-myeloid lineage bias and suggest the potential role of T cell epitopes in vaccine design and development.

## INTRODUCTION

The coronavirus disease 2019 (COVID-19) pandemic was caused by severe acute respiratory syndrome coronavirus 2 (SARS-CoV-2) infection. This highly infectious virus and its variants have had profound and devastating impact globally and are estimated to have caused more than 7 million deaths worldwide (https://covid19.who.int/). SARS-CoV-2 is a positive-sense RNA virus belonging to the coronavirus family. Its genome is ∼30 kb in size, contains 14 open reading frames (ORFs) and encodes 29 viral proteins. Virion structural proteins include the spike (S), nucleocapsid (N), envelope (E), and membrane (M) proteins. The S protein is comprised of two subunits (S1 and S2), where the S1 binds to the angiotensin-converting enzyme 2 (ACE2) receptor and S2 fuses with the cell membrane^1–3^. Many SARS-CoV-2 vaccines including the mRNA vaccines mRNA-1273 and BNT162b2, the viral vector-based vaccines Ad26.COV2.S, Sputnik V and AZD1222, and the protein-based vaccines NVX-CoV2373 and EpiVacCorona^4–11^ target S protein and prevent viral infection by inducing neutralizing antibodies that bind to that protein and inhibit virus entry into cells. Effectiveness of mRNA-1273-, BNT162b2- and Sputnik V-mediated protection against the original strain exceeds 90%, a value superior to other vaccines. S protein mutations can lead to emergence of variants of concern (VOCs). Since emergence of the original SARS-CoV-2 strain in December of 2019, Alpha (B.1.1.7), Beta (B.1.351), Gamma (P.1), Delta (B.1.617.2), and Omicron (B.1.1.529 and sublineages) have been defined as VOCs by the World Health Organization (WHO). Emergence of these VOCs can alter viral transmissibility, immune evasion, or vaccine effectiveness^4,5,9,12–15^. Therefore, it is necessary to design novel vaccines to ensure durable immune responses against them.

The immune system consists of innate and adaptive immune responses, the latter is key to controlling most viral infections and thus to vaccine effectiveness. The adaptive immune system is comprised of B, CD4^+^ T, and CD8^+^ T cells. Viral infection induces antigen-specific antibodies and generation of CD4^+^ and CD8^+^ T cells^16,17^. Antigen-specific CD4^+^ T cells then differentiate into a range of helper and effector cell types, like Th1 cells, which exhibit antiviral activities via producing IFN-γ and related cytokines, and T follicular helper cells (Tfh), which are critical for development of neutralizing antibody responses, memory B cells and long-term humoral immunity. CD4^+^ T cells signaling can activate CD8^+^ T cells, recruit innate cells and facilitate tissue repair. Antigen-specific CD8^+^ T cells are capable of killing virus-infected cells^16,18–23^. SARS-CoV-2 infection results in asymptomatic, clinically mild, or severe disease requiring hospitalization, and SARS-CoV-2 specific T cell and antibody responses are significantly associated with mild cases, suggesting that T cell responses are critical to control viral infection^18,24,25^. Previous analyses of SARS-CoV-2-specific CD4^+^ and CD8^+^ T cells responses in COVID-19 cases showed that not only S protein but also M, N, and nonstructural protein (NSP) stimulate T cell responses^19,26–29^. Moreover, last year, Arieta et al. reported that the T-cell-directed vaccine BNT162b4, which includes non-spike antigens, protects animals from severe SARS-CoV-2 infection^30^.

By contrast, trained immunity is defined as long-term adaptation of the innate immune system after initial contact with certain microbes, enabling faster, more robust responses to secondary challenge with homologous or heterologous pathogens. Trained immunity, which is promoted in part by changes in hematopoietic stem cell (HSC) differentiation, has recently emerged as a pivotal field of research in host defense against infections and vaccinations, as it confers broad protection against different infections and certain cancers^31–35^. Adding new antigens to vaccine design may generate trained immunity to provide broader and more durable immune responses compared with traditional spike-based vaccines.

Here, we asked whether introducing new antigens to induce anti-viral responses would enhance vaccine effectiveness and decrease disease risk due to escape variants. To do so, we designed a new mRNA vaccine consisting of the SARS-CoV-2 receptor-binding domain (RBD) of the spike protein, which is the smallest domain that directly binds to the ACE2 receptor, and new antigen epitopes. Others have shown that RBD mRNA vaccines elicit robust protective immunity against SARS-CoV-2^36–41^. We selected 20 non-RBD SARS-CoV-2 epitopes and 4 HIV epitopes, all of which stimulate T cell responses, to combine separately with RBD^28,42–46^. After screening and in vivo immune analysis, we identified G1-C mRNA vaccine (containing LVGLMWLSYFIASFRLFARTRSMWSFNPETNIL peptide from the SARS-CoV-2 membrane protein) as providing broader, more efficient protection against infection than the RBD mRNA vaccine in mice, and more robust spike-specific antibody production and T cell immune responses. Yarmarkovich et al. identified this membrane peptide as an epitope and predicted its presentation on HLA class I and class II^42^. Induced antibodies showed superior protection against infection by ancestral virus (WA1/2020, the first SARS-CoV-2 isolate detected in the United States), Beta, Delta and Omicron strains and promoted expansion of germinal center B and plasma cell populations. Remarkably, immunization with the G1-C vaccine also significantly increased proportions of hematopoietic stem cells (HSCs), progenitor cells and lymphoid lineage cells, suggesting that the vaccine modulates HSC differentiation and the balance of myeloid and lymphoid lineage cells in bone marrow (BM), potentially impacting trained immunity. Single cell RNA-sequencing (scRNA-seq) of BM cells revealed that transcription factor effectors of TGF-β and Wnt signaling may mediate some of these changes. Overall, our findings suggest that the G1-C mRNA vaccine induces a stronger adaptive immune response and better protection against SARS-CoV-2 infection than RBD-based vaccines and may regulate BM immune cells production.

## RESULTS

### SARS-CoV-2 mRNA vaccine design strategy

To determine whether new epitopes enhance RBD mRNA vaccine effectiveness, we selected 20 SARS-CoV-2 epitopes and 4 HIV epitopes to induce T cell immune responses ^28,42–47^ for inclusion in the RBD mRNA vaccine (Fig 1A). Epitope sequences fell into six groups, each containing four peptides (Fig 1B). We then generated N1-methylpseudouridine (m1Ψ)-modified RBD mRNA plus 24 new mRNA constructs, each harboring the RBD domain plus one new epitope (Fig 1A). These constructs contained a C-terminal human IgE signal peptide (SP) to promote secretion, untranslated regions (UTRs) to enhance translation, and 5’-cap modification and a poly(A) tail to increase stability and translation efficiency^48,49^. We also performed codon optimization to increase protein expression. We then synthesized m1Ψ-modified mRNA by in vitro transcription (IVT) of a linearized DNA plasmid, transfected HEK293T cells with these mRNA constructs and analyzed expression of encoded proteins by western blotting. Different mRNA constructs yielded varying levels of protein expression, potentially due to differences in translation efficiency or sequence-dependent effects from the incorporated epitope tags (Supplementary Fig 1A). We formulated mRNA-lipid nanoparticles (LNPs) using D-Lin-MC3-DMA: 1, 2-distearoyl-sn-glycero-3-phosphocholine: cholesterol: DMG-PEG2000 at a molar ratio of 50:10:38.5:1.5. Dynamic light scattering (DLS) analysis of mRNA-LNP physical properties indicated that mRNA-LNPs exhibited mean particle size of 91.6 nm, with −3.64 mV zeta potential and a favorable polydispersity index (PDI) (Supplementary Fig 1B).

**Figure 1.**
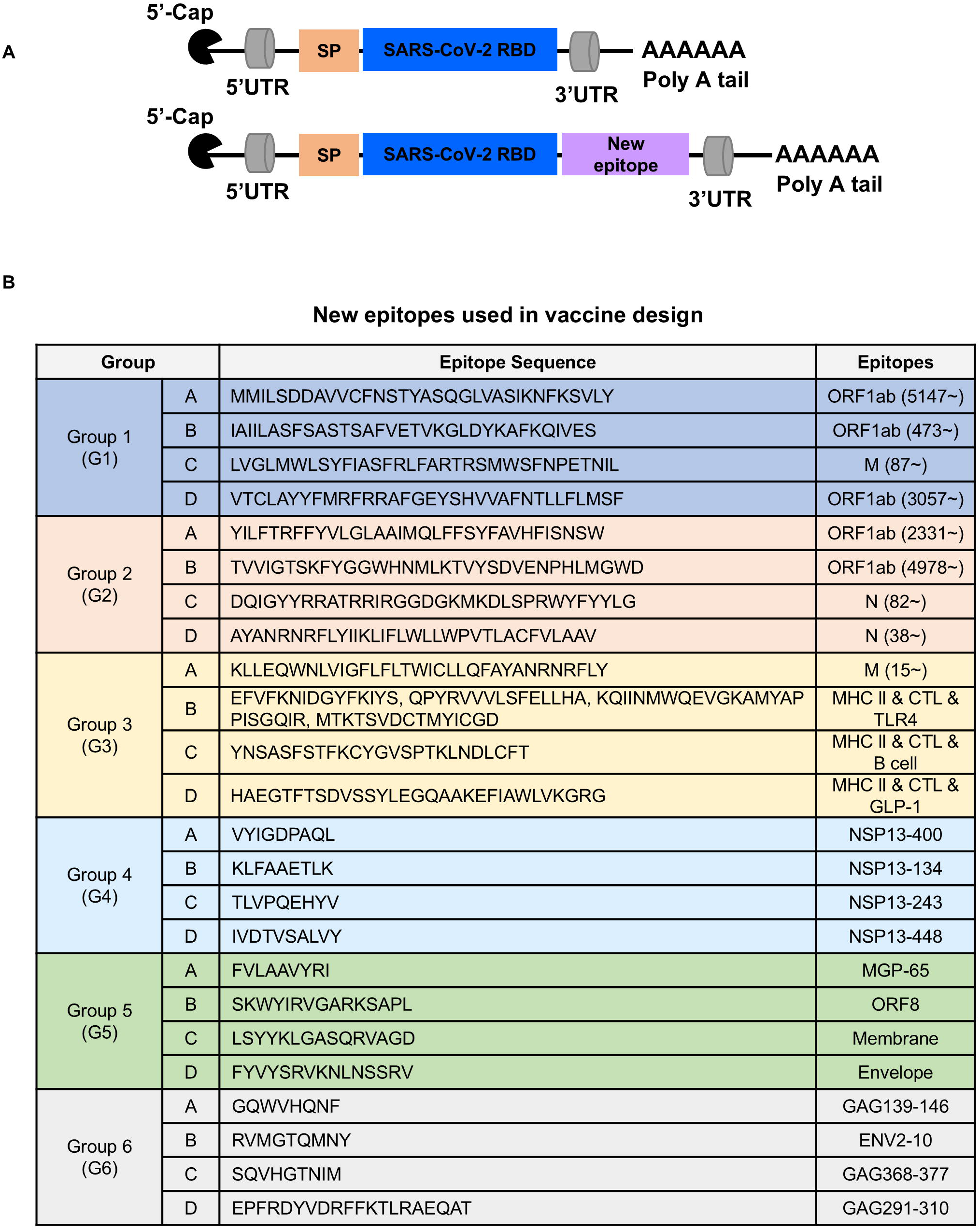
mRNA construct design strategy. (A) Schematic showing mRNA construct design. Constructs contain the N-terminal human IgE signal peptide (SP), 3’- and 5’- untranslated regions (UTRs), 5’-cap modification, and a poly-A tail, in addition to new epitopes. (B) Peptide sequences of all new epitopes tested in the study. Epitopes are divided into six groups each consisting of four unique peptides. Epitope origin is also shown.

### mRNA Vaccine Group 1 elicits robust antibody and T cell responses in mice

To evaluate immune responses induced by each of the 6 groups of candidate mRNA vaccines, we immunized C57BL/6J adult mice intramuscularly twice, 2 weeks apart, with vaccine pools (10 μg) containing 4 peptides from each of the 6 groups on days 0 and 14. We then collected blood and spleen 2 weeks later to analyze immune responses (Fig 2A). Using GFP mRNA and RBD mRNAs as respective negative and positive controls, we analyzed both total and anti-RBD IgG, IgA and IgM antibody responses. We observed no significant differences in GFP-, RBD- or Groups 1-6-induced total IgG, IgA and IgM levels (left graph of panels Fig 2B, C, D). However, analysis of RBD-specific IgG, IgA, and IgM revealed that both the RBD group and Groups 1-6 triggered RBD-specific antibody responses. Notably, Groups 2-6 exhibited antibody levels comparable to those generated by the RBD vaccine, while only the Group 1 vaccine significantly induced a more robust RBD-specific IgG, IgA, and IgM antibody response than did the RBD vaccine (right graph, Fig 2B, C, D). We also compared the levels of WA1 RBD-specific IgG and XBB.1.5 RBD-specific IgG induced by 5 μg Moderna mRNA vaccine (2023–2024 formula, encoding the Spike glycoprotein of the SARS-CoV-2 Omicron variant lineage XBB.1.5) and 10 μg RBD mRNA vaccine. We observed that the XBB.1.5 RBD–specific IgG levels induced by the two vaccines were generally within a similar range, whereas the Moderna vaccine produced lower WA1 RBD–specific IgG responses. These findings indicate that the RBD vaccine can elicit a robust immune response under the tested conditions. (Supplementary Fig 2A).

**Figure 2.**
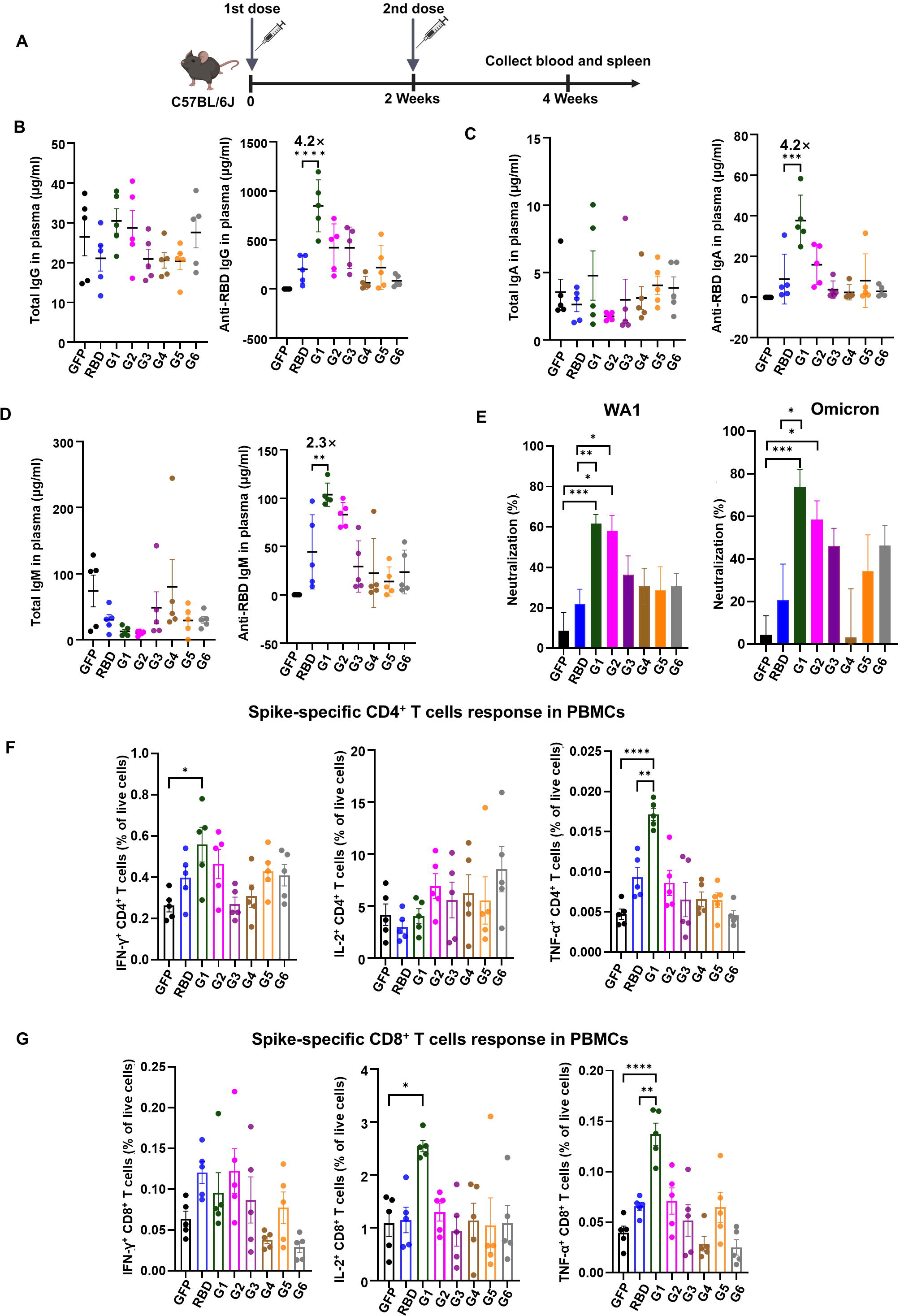
Screening of mRNA vaccine candidates by analysis of antibody and memory T cell responses. (A) Immunization procedure for mRNA vaccine candidates in C57BL/6J mice. Mice (n=5/group) were vaccinated with two doses (10 μg/dose) of mRNA-LNPs at 2-week intervals. Blood was collected for antibody and T cell response analyses. (B-D) IgG, IgM, and IgA titers in plasma of immunized mice, as quantified by ELISA. (E) Neutralizing activity of immunized mice plasma against WA1 and Omicron SARS-CoV-2 pseudoviruses. Mouse plasma was diluted 2500-fold for the assay (n=5). (F, G) Spike-specific CD4^+^ and CD8^+^ T cell responses induced by mRNA vaccines in PBMCs, as measured by intracellular staining of IFN-γ^+^, IL-2^+^ and TNFα^+^ after stimulation with the spike peptide pool. Asterisks represent statistical significance (*p < 0.05; **p < 0.01; ***p < 0.001; ****p < 0.0001) as determined by one-way ANOVA (B, C, D, F, G) and unpaired t-test (E). Error bars indicate standard error of the mean (SEM).

We next performed a SARS-CoV-2 pseudovirus neutralization assay to analyze neutralization activity of antibodies induced by different vaccines. RBD and Group 1-6 vaccines exhibited 21.90%, 61.64%, 58.15%, 36.28%, 30.57%, 28.66%, and 30.68% neutralizing activity, respectively, against USA-WA1/2020 (WA1) pseudovirus, and 20.58%, 73.73%, 58.53, 46.08%, 3.07%, 34.36%, and 46.22% neutralizing activity against Omicron pseudovirus after 2500-fold dilution of mouse plasma. Compared with the RBD vaccine, Group 1 had better neutralization activity against WA1 and Omicron pseudovirus (Fig 2E). Neutralizing ability against Beta and Delta SARS-CoV-2 pseudovirus was also greater for Group 1 relative to RBD (Supplementary Fig 2B).

Next, we analyzed T cell immune responses in peripheral blood mononuclear cells (PBMCs) and spleen. To do so, we analyzed spike-specific T cell responses in PBMCs stimulated 16h with spike peptide pools based on levels of tumor necrosis factor (TNFα), interferon-γ (IFN-γ), and interleukin-2 (IL-2) ^50–53^ and found that the Group1 vaccine increased numbers of IFN-γ^+^ CD4^+^ T, TNFα^+^ CD4^+^ T, IL-2^+^ CD8^+^ T, and TNFα^+^ CD8^+^ T cells relative to GFP controls (Fig 2F, G, gating strategy and representative plots are shown in Supplementary Fig 9). In spleen, the Group1 vaccine increased proportions of IL-2^+^ CD4^+^ T, TNFα^+^ CD4^+^ T, IL-2^+^ CD8^+^ T, and TNFα^+^ CD8^+^ T cells relative to the RBD control (Supplementary Fig 2C, D). Overall, these results suggest that the Group 1 vaccine significantly induces spike-specific IgG, IgA, and IgM production and can provide robust neutralization of WA1, Beta, Delta, and Omicron pseudoviruses. The vaccine also significantly increased spike-specific CD4 and CD8 T cell activation in PBMCs and spleen relative to GFP and RBD controls. These data overall indicate that the Group 1 mRNA vaccine generates antibody and T cell immune responses superior to the RBD vaccine. As noted above, different mRNA constructs express proteins at varying levels (Supplementary Fig 1A), which may influence vaccine-induced immune responses. This variation limits our ability to attribute immunogenicity differences solely to epitope design.

Importantly, G1 exhibits a striking exception: despite relatively low protein expression by Western blot, it induces strong immune responses that match or exceed those of higher-expressing constructs. This dissociation between expression level and immunogenicity provides strong evidence for a design-driven effect independent of protein abundance.

### The G1-C mRNA vaccine elicits potent antibody, T cell and B cells responses in mice

Next, we asked which construct in the pool contributes most to Group 1 vaccine-induced adaptive immune responses using vaccines consisting of single mRNA constructs. To do so, we immunized C57BL/6J mice intramuscularly with 10 μg vaccine on days 0 and 14 and collected blood samples, spleen and BM 2 weeks later (Fig 3A). Consistent with our preliminary screen, antibody production, neutralizing activity, and spike-specific T cell responses following immunization with each of the 4 Group 1 mRNA vaccines was comparable to changes shown above in Fig 2 (Fig 3B, D, E, Supplementary Fig 3A). Analysis of vaccination-induced IgG antibody responses confirmed that Group 1 induces higher levels of RBD-specific IgG than the RBD vaccine (Fig 3B). Notably, among the 4 vaccines, G1-C induced WA1 RBD-specific IgG levels that were 8.2-fold higher and XBB.1.5 RBD-specific IgG levels that were 16.6-fold higher than those induced by the RBD vaccine (Fig 3B). When we evaluated neutralizing activity in plasma from RBD and G1-C immunized mice, both showed concentration-dependent inhibition of WA1 and Omicron pseudovirus infection. The IC50 (half-maximum inhibitory concentration) of plasma neutralization titers from mice vaccinated with RBD or G1-C vaccine were 1213 or 2149 against WA1 pseudovirus, respectively, and the IC50 against omicron was 681 and 2386, respectively (Fig 3C). We also performed a neutralization assay of WA1, beta, delta, and omicron pseudoviruses using 2500-fold diluted plasma and found that G1-C vaccine exhibited the highest neutralizing activity among 4 Group 1 mRNA vaccines, and that activity was significantly better than that of RBD vaccine (Supplementary Fig 3A).

**Figure 3.**
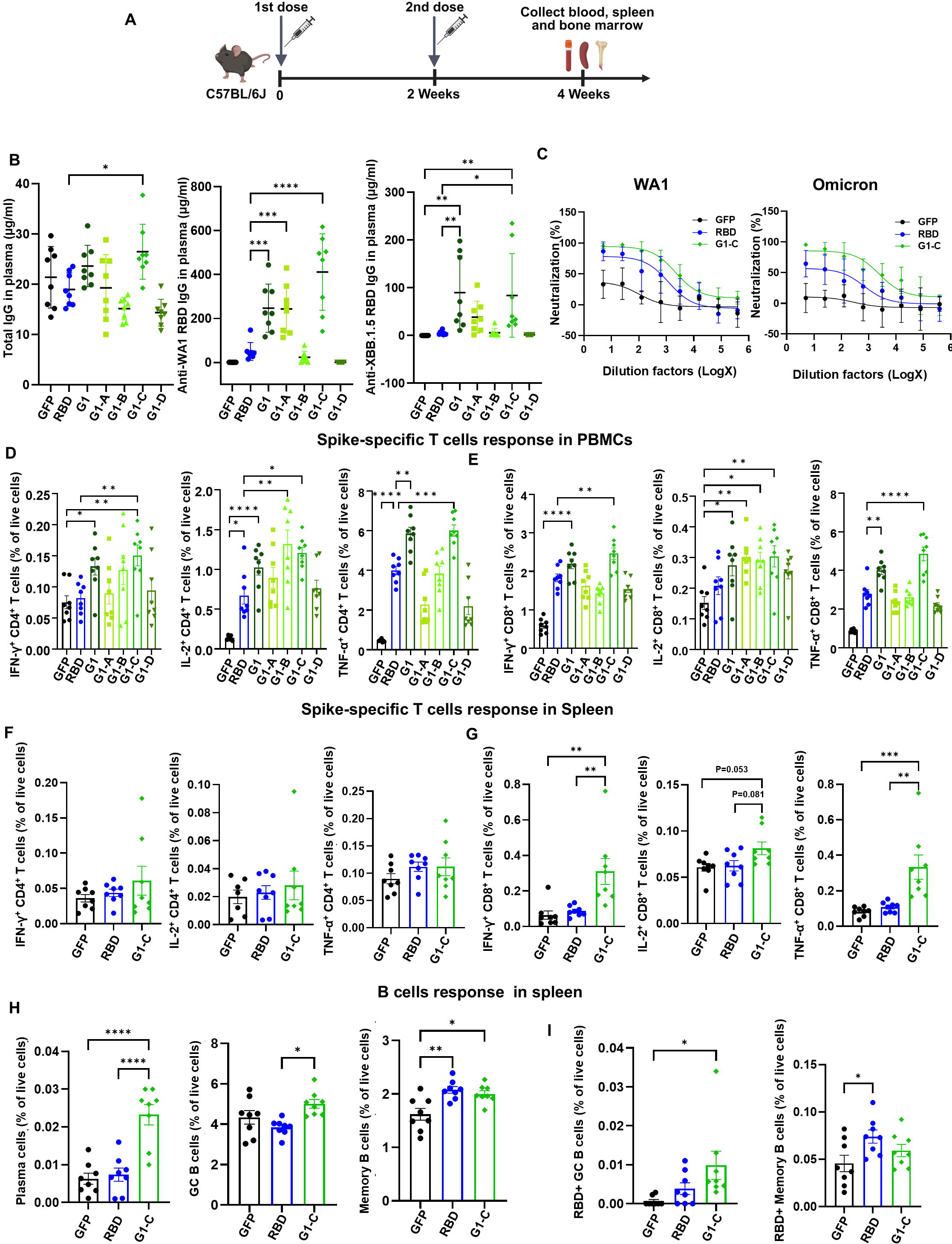
The G1-C mRNA vaccine induces robust antibody and T and B cell responses. (A) Immunization procedure for mRNA vaccine candidates in C57BL/6J mice. Mice (n=8/group) were vaccinated with two doses (10 μg/dose) of mRNA-LNPs at 2-week intervals. Blood, spleen, and BM were harvested to assess antibody responses and adaptive immune responses. (B) Total IgG, WA1 RBD-specific IgG and XBB.1.5 RBD-specific IgG titers in plasma, as detected by ELISA. (C) Neutralization activity of immunized mice plasma against WA1 and omicron SARS-CoV-2 pseudoviruses. Serially-diluted plasma was used for the assay (n=6). (D-G) Spike-specific CD4^+^ and CD8^+^ T cell responses induced by mRNA vaccines in PBMCs (D, E) and spleen (F, G), based on intracellular staining of IFN-γ^+^, IL-2^+^ and TNFα^+^ after stimulation with spike peptide pools. (H) The proportion of plasma (Blimp-1^+^CD138^+^), germinal center B (B220^+^Fas^+^CD38^low^) and memory B cells (B220^+^CD80^+^CD21^+^) in spleen after vaccination. (I) The proportion of RBD-binding germinal center B and memory B cells in spleen after vaccination. Asterisks represent statistical significance (*p < 0.05; **p < 0.01; ***p < 0.001; ****p < 0.0001) as determined by one-way ANOVA. Error bars indicate standard error of the mean (SEM).

We also analyzed TNFa-, IL-2-, and IFNg-positive T cell populations as shown above in Fig 2 and found that G1-C significantly increased spike-specific CD4 and CD8 T cell responses in PBMCs compared to GFP control and RBD vaccine (Fig 3D, E) and spike-specific CD8 T cell responses in spleen (Fig 3G), but did not augment spike-specific CD4 T cell responses in spleen (Fig 3F). Since the novel G1-C epitope is from the SARS-CoV-2 membrane protein, we also analyzed membrane-specific T cell responses. As expected, G1-C significantly increased levels of M-specific CD4 and CD8 T cell activation relative to the RBD group (Supplementary Fig 3B, C). To further characterize the CD8 T cell responses, we analyzed cytotoxic activity and CD8 T cell subsets in spleen. We found that G1-C vaccine significantly increased Granzyme B^+^ CD8 T cells and effector CD8 T cells, while decreased naïve CD8 T cells compared to GFP control (Supplementary Fig 4A, gating strategy is shown in Supplementary Fig 12A). These results indicate that G1-C vaccine promotes CD8 T cell activation, and differentiation. Germinal centers serve as a hub for differentiation of long-lived plasma cells and mature memory B cells, which are required for sustained antibody production and long-term immune memory^54–56^. Therefore, we assessed the dynamics of germinal center (GC) cells. Analysis of splenic B cell responses showed that the G1-C vaccine strongly increased the number of GC B and plasma cells compared to RBD vaccine and RBD-specific GC B cell populations compared to GFP control (Fig 3H, I, gating strategy and representative plots are shown in Supplementary Fig 10). Additionally, we also observed an increase of classical dendritic cells (cDCs) in G1-C compared with GFP and RBD groups (Supplementary Fig 4B, gating strategy is shown in Supplementary Fig 12B). The increase of cDCs indicates enhanced antigen-presenting capacity, which likely supports the activation of T cell responses and B cell responses. Overall, these findings suggest that the G1-C mRNA vaccine may produce broad, long-term immune memory and protection against SARS-CoV-2.

### The G1-C mRNA vaccine regulates HSC differentiation in bone marrow

Trained immunity emerges via direct recognition of Microbe-Associated Molecular Patterns (MAMPs) by Pattern Recognition Receptors (PRRs) or by cytokines released during induction of the host response^31^. Moreover, BM hematopoietic stem and progenitor cells (HSPCs) can be reprogrammed to boost myeloid and lymphoid cell production during infection by a process termed emergency myelopoiesis and lymphopoiesis^32–34^. To determine whether such reprogramming occurs in response to the G1-C mRNA vaccine, we performed flow cytometry to assess HSPCs from vaccinated mice. We observed that the G1-C mRNA vaccine significantly increased proportions of long-term hematopoietic stem cells (LT-HSCs), short-term hematopoietic stem cells (ST-HSCs), common myeloid progenitors (CMPs), and lymphoid-primed multipotent progenitors (LMPPs) relative to the RBD vaccine (Fig 4A, B, gating strategy and representative plots are shown in Supplementary Fig 11A, B). We then performed single-cell RNA sequencing (scRNA-Seq) of 9 samples from GFP, RBD and G1-C groups to assess differential gene expression. Analysis was performed in a total of 115388 single cells, and clusters were annotated by reference mapping and visualized by Uniform Manifold Approximation and Projection (UMAP)^57^ (Fig 4C). Analysis of 15 cell types revealed that vaccination significantly altered the proportions of LMPPs, B cells, pre-B cells, neutrophils, and NK cells (Fig 4D). Quantification of absolute cell numbers validated these changes, demonstrating significant increases in lymphoid populations following G1-C vaccination (Supplementary Table 2). Cell type specific markers, including *Ebf1*, *Ly6d*, *Cd3d*, *Gzma*, *Lcn2*, and others, were evaluated and visualized on UMAP (Fig 4E, Supplementary Fig 5B), and key markers for each cluster were visualized in a dot plot (Supplementary Fig 5A).

**Figure 4.**
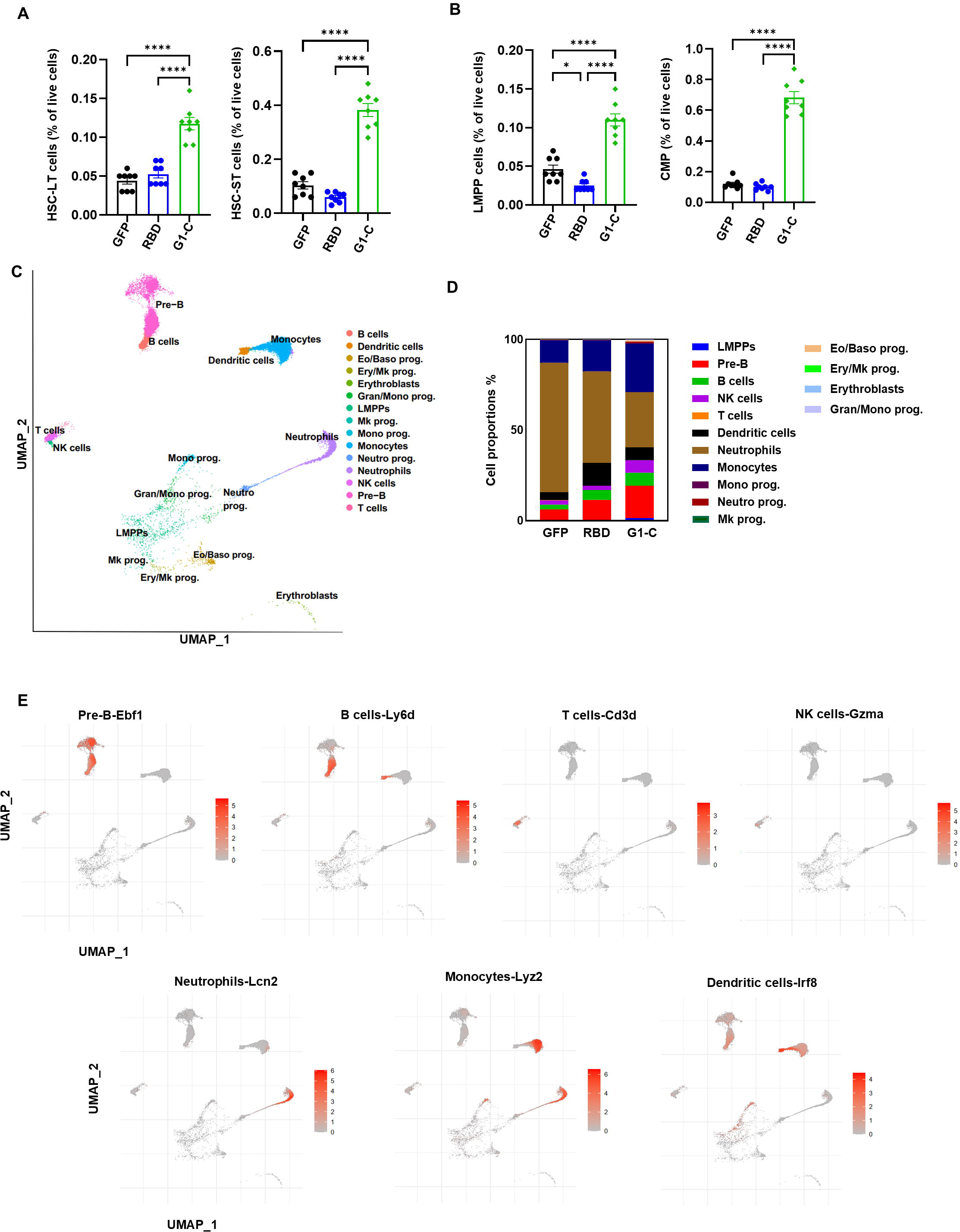
G1-C mRNA vaccine immunization modulates hematopoietic stem cell development in bone marrow. (A) The proportion of long-term hematopoietic stem cells (LT-HSCs) and short-term hematopoietic stem cells (ST-HSCs) in BM after vaccination (n=8). LT-HSCs (Lin^-^Sca1^+^c-Kit^+^CD34^-^CD135^-^) and ST-HSCs (Lin^-^Sca1^+^c-Kit^+^CD34^+^CD135^-^) were gated after removing doublets and dead cells. (B) The proportion of common myeloid progenitors (CMPs) and lymphoid-primed multipotent progenitors (LMPPs) in BM after vaccination (n=8). CMPs (Lin^-^Sca1^-^c-Kit^+^CD34^+^CD16/32^-^) and LMPPs (Lin^-^Sca1^+^c-Kit^+^CD34^+^CD135^+^) were gated after removing doublets and dead cells. (C) UMAP for scRNA-seq data from BM showing identified 15 cell types. Single cells are color-coded by cluster annotation (n = 3 /group). (D) Bar graph showing proportions of 15 cell types in GFP, RBD and G1-C vaccine immunized BM samples as identified by scRNA-seq. (E) Expression of marker genes for pre-B, B, T, and NK cells, and neutrophils, monocytes and dendritic cells, highlighted on UMAP. BM samples were collected after two doses of mRNA-LNPs administered at 2-week intervals. Asterisks represent statistical significance (*p < 0.05; **p < 0.01; ***p < 0.001; ****p < 0.0001) as determined by one-way ANOVA. Error bars indicate standard error of the mean (SEM).

The balance of lymphoid and myeloid lineage cells differentiated from BM HSCs is critical to immune function. Thus we assessed effects of G1-C vaccination on these populations by analyzing proportions of lymphoid (NK, B and T cells) and myeloid (neutrophils, monocytes and dendritic cells) lineage cells separately. Interestingly, G1-C vaccination increased the proportion of lymphoid lineage cells and decreased that of myeloid lineage cells relative to GFP and RBD controls (Fig 5A, C). Consistent with our scRNA-seq data, flow cytometry analysis also showed that G1-C vaccination significantly increased the proportion of NK cells, B cells and monocytes, and decreased neutrophil levels in BM (Fig 5B, D, gating strategy is shown in Supplementary Fig 11C).

**Figure 5.**
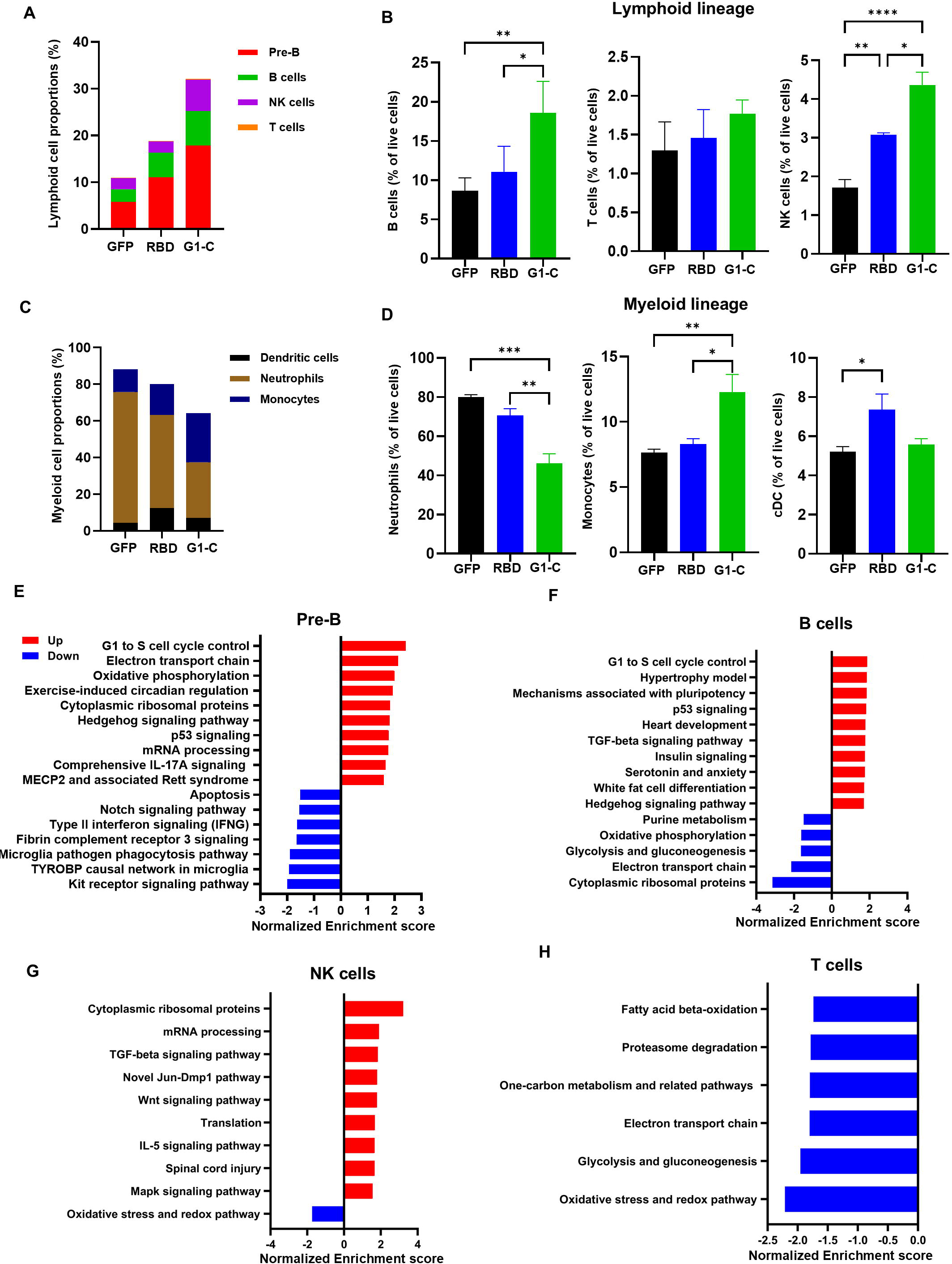
Immunization with the G1-C mRNA vaccine promotes differentiation of lymphoid lineage cells. (A) Bar graph showing proportions of lymphoid lineage cells after immunization with indicated vaccines, based on scRNA-seq. (B) Analysis of the proportion of B, T and NK cells in BM of mice immunized with indicated vaccines, based on flow cytometry (n=4). (C) Bar graph showing proportions myeloid lineage cells in BM of mice immunized with indicated vaccines, based on scRNA-seq. (D) Analysis of proportion of neutrophils, monocytes and dendritic cells in BM of mice immunized with indicated vaccines, based on flow cytometry (n=4). (E-H) Top 10 up- and down-regulated canonical pathways in pre-B, B, T and NK cells. DEGs were ranked using the DESeq2 STAT column, and fGSEA was run using both Wikipathways and REACTOME databases. Enriched pathways were filtered using an FDR cutoff of 10%. BM samples were collected after two doses of mRNA-LNPs administered at 2-week intervals. Asterisks represent statistical significance (*p < 0.05; **p < 0.01; ***p < 0.001; ****p < 0.0001) as determined by one-way ANOVA. Error bars indicate standard error of the mean (SEM).

To identify mechanisms potentially underlying these effects, we performed enrichment analysis of canonical signaling pathways after G1-C or RBD vaccination and identified the top 10 up- or down-regulated pathways in lymphoid lineage cells (Fig 5, E-H). TGF-β signaling, which regulates B and NK cell development and function ^58–61^, was upregulated in both of those cell types. Wnt signaling, which promotes NK, T and B cell development^62,63^, was also upregulated in NK cells. Pathways controlling the G1 to S cell cycle transition, which is crucial to control eukaryotic cell proliferation^64–67^, were upregulated in Pre-B and B cells (Fig 5 E-H). Relevant to this finding, immunization with the G1-C vaccine promoted the G1 phase to S phase transition (Fig 6A), an outcome that may promote B cell proliferation.

**Figure 6.**
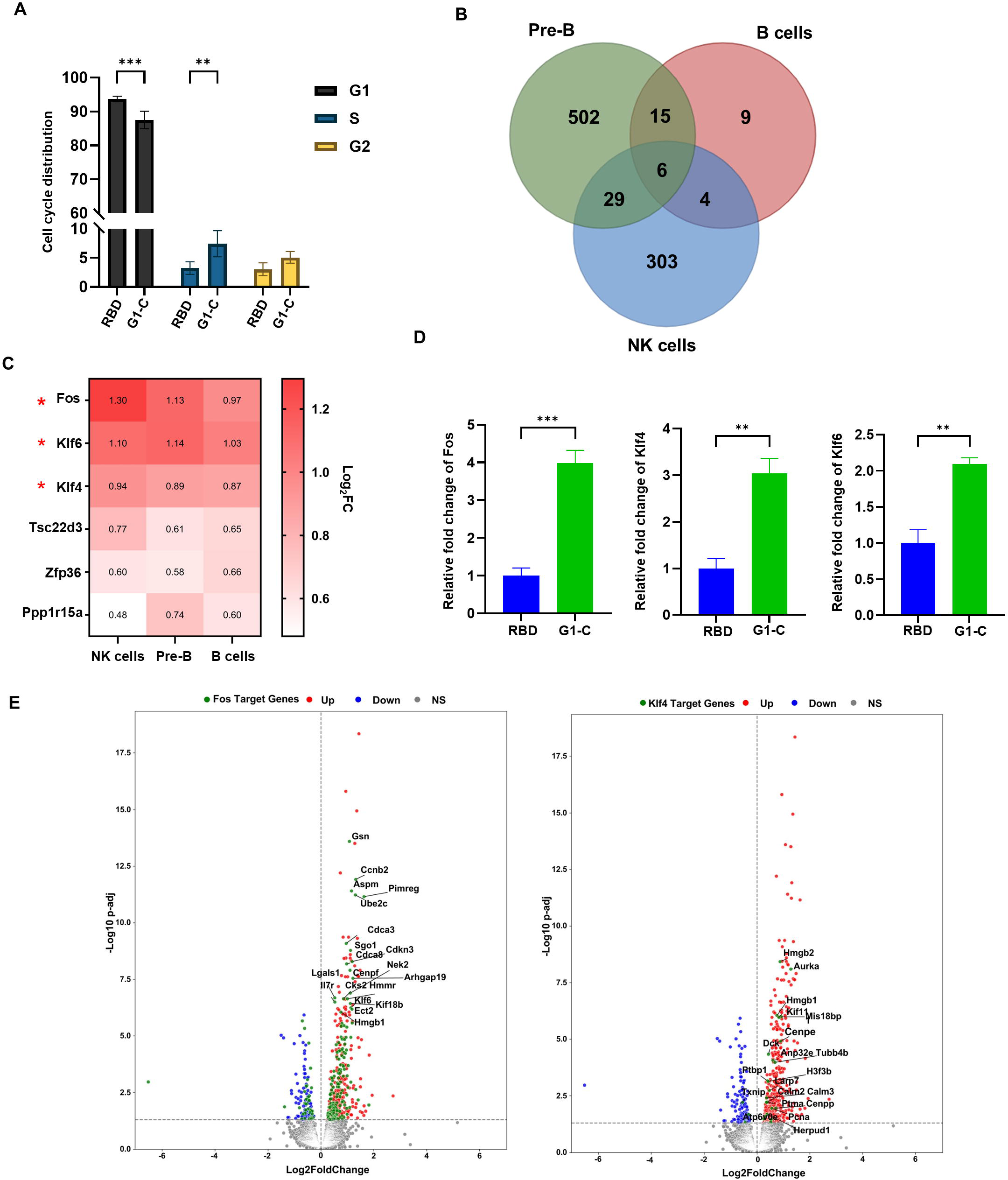
Immunization with G1-C mRNA vaccine modulates B and NK cell function in bone marrow. (A) Cell cycle distribution in B220^+^ B cells (n=4) following immunization of mice with RBD or G1-C mRNA vaccines. FxCycle™ Violet stain was used to assess DNA content. (B) Venn plot of DEGs in pre-B, B and NK cells. DEGs were identified in cells from G1-C-versus RBD-immunized mice using DESeq2. DEGs with padj<0.05 were used for overlap analysis. (C) Heatmap of 6 overlapped DEGs in analysis shown in (B). log2FC values of G1-C versus RBD are indicated in red boxes. (D) RT-PCR analysis of fold-changes in expression of Fos, Klf4, and Klf6 in BM cells (5×10^6^) from mice immunized with G1-C versus RBD vaccines. Values are normalized to Gapdh; shown are relative fold-changes (n=4). (E) Volcano plots showing differential expression of Fos (left) and Klf4 (right) target genes in pre-B cells from mice immunized with G1-C versus RBD vaccine, as analyzed using DESeq2. ChIP-Atlas 3.0 was used to identify Fos and Klf4 target genes at the TSS ± 5k from M. musculus (mm10). Red dots, upregulated genes; blue, downregulated genes; gray, genes not significantly changed. Green dots indicate DEGs overlapping in both Fos or Klf4 target genes. BM samples were collected after two doses of mRNA-LNPs administered at 2-week intervals. Asterisks represent statistical significance (*p < 0.05; **p < 0.01; ***p < 0.001; ****p < 0.0001) as determined by two-way ANOVA (A) or one-way ANOVA (D). Error bars indicate standard error of the mean (SEM).

We next analyzed differentially expressed genes (DEGs) in lymphoid lineage cells including B cells, pre-B cells, T cells and NK cells, comparing the G1-C and RBD. We identified 6 overlapping upregulated genes in B cells, pre-B cells, and NK cells: *Fos*, *Klf4*, *Klf6*, *Tsc22d3*, *Zfp36*, and *Ppp1r15a*. Among these, transcript levels of *Fos*, *Klf4*, and *Klf6* were increased by approximately 2-fold across all three cell types. (Fig 6B, C). The changes of *Fos, Klf4*, and *Klf6* mRNA expression levels were verified in RBD and G1-C immunized mice BM cells by RT-qPCR (Fig 6D).

From scRNA-seq data, we found that TGF-β and Wnt signaling pathways were activated in G1-C compared to RBD (Fig 5F, G). To further confirm the activation of TGF-β and Wnt signaling pathways, the expression levels of signal pathways related genes were analyzed in RBD and G1-C immunized mice BM cells. The RT-qPCR results showed that *Fos, Smad1, Prkcb* and *Prkcq* were significantly upregulated in G1-C group, in agreement with sequencing data (Fig 6D, Supplementary Fig 6A).

To validate pathway involvement, we used pharmacological modulators of TGF-β and Wnt/β-catenin signaling. KRFK TFA, a thrombospondin-1 (TSP-1)-derived peptide, activates TGF-β signaling^68,69^, while CJJ300 inhibits this pathway by disrupting the TGF-β-TβR-I-TβR-II signaling complex ^70^. Similarly, BIO activates Wnt/β-catenin signaling through GSK3 inhibition^71,72^, while MSAB inhibits this pathway by promoting β-catenin degradation ^73^. Thus, to activate either of these signaling pathways, we isolated BM cells from C57BL/6J mice, treated them 48 hrs with KRFK TFA (100 μM) or BIO (1 μM), and analyzed proportions of T, B and NK cells by flow cytometry. KRFK TFA treatment significantly increased the proportion of all three cell types, while BIO treatment significantly increased the proportion of B and NK cells (Supplementary Fig 6B, C). To inhibit TGF-β or Wnt/β-catenin signaling pathways, we treated BM cells 48 hrs with CJJ300 (10 μM) or MSAB (4 μM) and analyzed proportions of T, B and NK cells by flow cytometry. Both treatments significantly increased the proportion of B and NK cells (Supplementary Fig 6D, E). Collectively, these pharmacological validation experiments demonstrate that G1-C mRNA vaccination promotes lymphoid cell differentiation and development in bone marrow through regulation of TGF-β and Wnt/β-catenin signaling pathways.

We then used the ChIP-Atlas 3.0. to identify target genes of Fos, Klf4 and Klf6 in pre-B cells, B cells and NK cells. In order to do that, we overlapped the transcription factors targeted genes and DEGs (G1-C vs RBD) in these 3 cell types. DEGs (G1-C vs RBD) in different cell types were visualized using volcano plots. Overlapping genes were marked in green (Fig 6E, Supplementary Fig 6). Interestingly, multiple Fos- and Klf4-targeted genes were present among DEGs identified in our analysis (Fig 6E, Supplementary Fig 7). These data suggested that Fos, Klf4 and Klf6 are key transcription factors that regulate development of lymphoid lineage cells in G1-C immunized BM.

### The G1-C mRNA vaccine provides potent protection against SARS-CoV-2 in K18-hACE2 mice

We performed sequence alignment across different SARS-CoV-2 variants and found that G1-C epitope is highly conserved (Supplementary Fig 8). We next compared protective efficacy of G1-C and RBD vaccines in K18-hACE2 transgenic mice. To do so, we immunized mice intramuscularly (IM) with two doses of 10 μg vaccine, and then, 2 weeks after booster, intranasally (i.n.) inoculated mice with 1×10^4^ plaque-forming units (PFU) of D614G SARS-CoV-2 virus (Fig 7A). Mice were monitored for body weight and clinical scores daily post-infection (p.i.) until sacrifice. Body weight decreased by 10.2%, 3.2%, and 0.2% in GFP, RBD, and G1-C vaccinated mice, respectively (Fig 7B). Moreover, clinical scores of GFP, RBD, and G1-C immunized mice were 4 (sick, very ruffled coat, slightly closed/inset eyes, walking but no scurrying, mildly lethargic), 3 (ruffled coat over the entire body, active, alert), and 2 (slightly ruffled coat around head and neck, active, alert), respectively (Fig 7C), suggesting overall that both RBD and G1-C vaccination protect mice against severe SARS-CoV-2 infection, while G1-C shows more protective effects.

**Figure 7.**
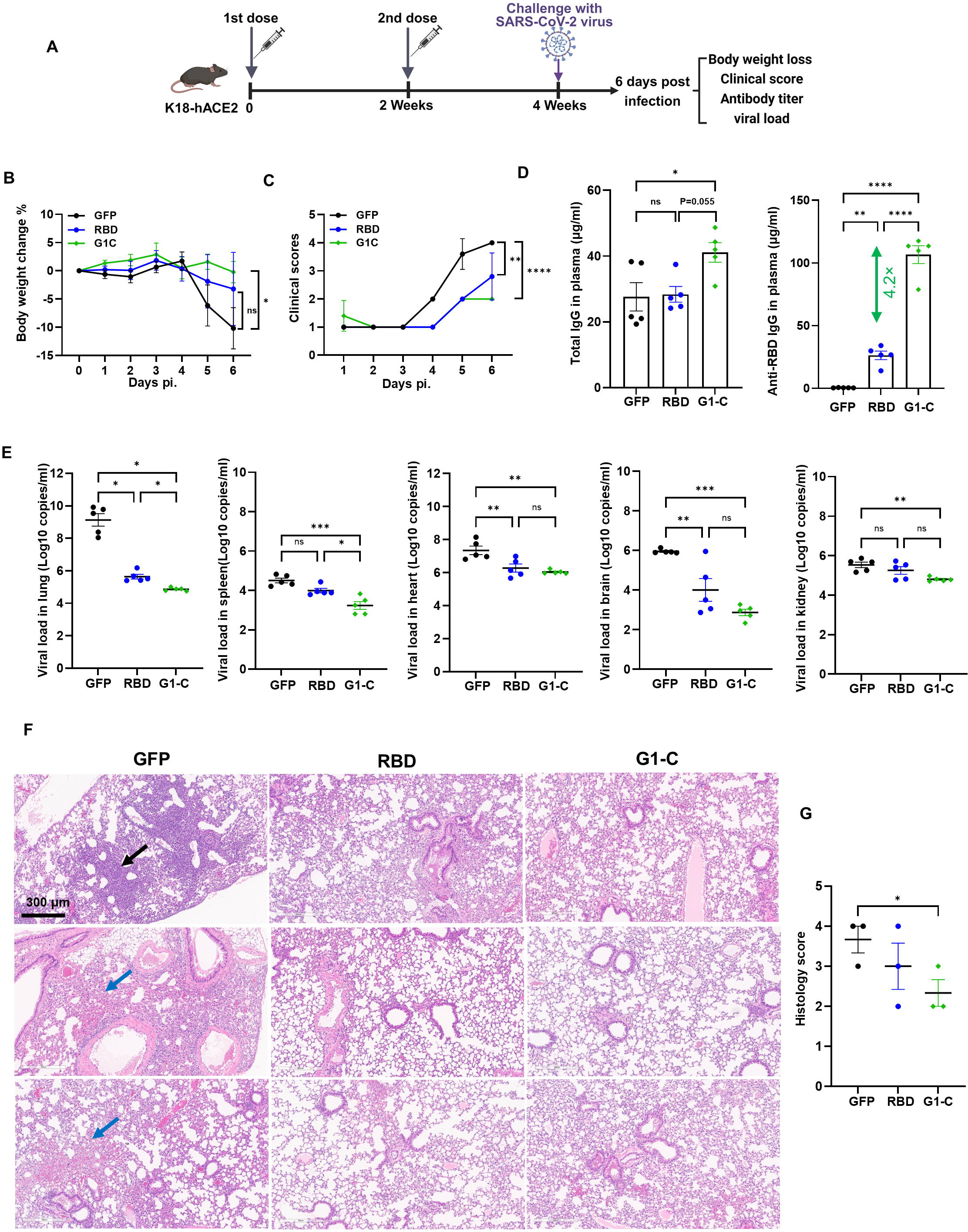
The G1-C vaccine provides better protection against SARS-CoV-2 infection than the RBD vaccine. (A) Immunization and virus challenge procedures used to immunize K18-hACE2 mice with RBD, G1-C mRNA and control (GFP) vaccines. Mice (n=5/group) received two doses (10 μg/dose) of mRNA-LNPs at days 0 and days 14 and then were challenged with 1×10^4^ PFU D614G SARS-CoV-2 at days 28. Body weight, scores, antibody responses, viral load and B cell responses were analyzed in all three groups. (B) Changes in body weight were monitored in indicated groups after virus challenge until euthanasia. (C) Clinical scores, ranging from low (mild disease) to high (more severe disease), were monitored in indicated groups after virus challenge until euthanasia. (D) Total IgG- (left) and RBD-specific (right) IgG titers in plasma of indicated mice, as detected by ELISA. (E) Viral RNA loads in lung, spleen, heart, brain, and kidney in indicated mice, as detected by RT-qPCR. (F) Evaluation pathological changes in lungs of indicated mice based on hematoxylin and eosin (H&E) staining (scale bar, 300 μm). Black arrow, inflammation; blue arrows, pulmonary hemorrhage. (G) Histopathological score of lung tissue from infected animals 6 days post-infection. Asterisks represent statistical significance (*p < 0.05; **p < 0.01; ***p < 0.001; ****p < 0.0001) as determined by one-way ANOVA. Error bars indicate standard error of the mean (SEM).

We then measured antibody titer, viral load, and tissue pathology in lungs at harvest (Fig 7A) in all three groups. Analysis of total IgG and anti-RBD IgG showed that G1-C-immunized mice produced more total IgG than GFP- or -RBD immunized mice and produced 4.2-fold more anti-RBD IgG than did RBD-immunized mice (Fig 7D). Analysis of viral titers in multiple tissues showed that RBD and G1-C vaccination significantly decreased viral load in lung, brain and heart relative to controls. Notably, G1-C immunized mice showed lower viral load than the RBD group in lung and spleen and showed lower viral load than the control group in kidney (Fig 7E). To further assess these pathological changes, we performed H&E staining of paraffin-embedded lung sections from immunized mice 6 days post-infection and observed that immunization with RBD or G1-C vaccine significantly decreased inflammation and hemorrhage likely caused by viral infection (Fig 7F, G). Overall, these data strongly suggest that G1-C mRNA vaccine provides better protection efficacy against D614G SARS-CoV-2 infection relative to the RBD vaccine.

## DISCUSSION

SARS-CoV-2 mRNA vaccines provide effective protection from viral infection and severe disease through a combination of immune responses, including neutralizing and non-neutralizing antibodies as well as CD4⁺ and CD8⁺ T cell responses. However, their protection efficacy against emerging variants is decreased and immune memory is relatively short lived. Such decreases in neutralizing antibody titers over time could lead to an increase in breakthrough infections^4,5^. Therefore, it is necessary to design new vaccines that provide durable immune responses against emerging variants. Immune responses by T cells are a critical part of adaptive immunity system. Upon recognizing viral antigen epitopes, antigen-specific T cells can differentiate into a range of helper and effector cell types to produce anti-viral cytokines, clear virus-infected cells or assist in B cell maturation^16–19^. Memory T cells in lung can be detected for at least 10 months after SARS-CoV-2 infection in human and the number of SARS-CoV-2-specific T cells in lungs correlates with effective clinical control, suggesting that memory T cells function to limit re-infection^18^. T cells recognize the spike protein, as well as other SARS-CoV-2 proteins, including nucleocapsid (N), membrane (M), and nonstructural protein (NSP) antigens^26,27^, indicating that these antigens could be considered in vaccine design to enhance protection efficacy and enable development of pan-coronavirus vaccines. Recently, the new T-cell-directed vaccine BNT162b4 was found to protect animals from severe disease and enhance viral clearance by combination with BNT162b2^30^. Analysis of the T cell immune response in mice spleen showed that BNT162b4 could enhance T cell immunity but did not affect spike-specific antibodies level indicating the critical role of T cell epitopes in vaccine design^30^.

We designed new mRNA vaccines consisting of SARS-CoV-2 RBD and new epitopes and then used them to immunize C57BL/6J mice and assess adaptive immune responses, among them, antibody titer, neutralization activity, and spike-specific T and B cell responses. One of our vaccines, the G1-C mRNA vaccine, induced higher levels of WA1 RBD-specific and XBB.1.5 RBD-specific antibodies, provided better and broader neutralization activity against WA1/2020, Beta, Delta and Omicron strains, and induced a better spike-specific T cell response than the RBD-based vaccine. We also observed expansion of populations of germinal center B and plasma cells in G1-C-vaccinated animals. These findings indicate that G1-C mRNA vaccine potentially provides a better capacity to resolve severe COVID19 cases than existing vaccines. When we evaluated protective efficacy of G1-C vaccine against D614G SARS-CoV-2 in K18-hACE2 transgenic mice, we found that relative to mice immunized with RBD vaccine, G1-C-immunized mice did not exhibit significant body weight loss and their viral titers were significantly reduced in multiple tissues.

The bacillus Calmette-Guérin (BCG) tuberculosis vaccine can induce training immune responses and provide protection against non-related to tuberculosis viral infections^31^. Trained immunity enhances immune responses against different infections by reprogramming HSCs and immune cells and altering cytokine production^33,74^. In our study, we found that the G1-C mRNA vaccine reprograms BM HSCs, lymphoid and myeloid lineage cells. Compared with RBD vaccine, G1-C vaccine significantly increased the number of HSCs, progenitors, B cells and NK cells. The G1-C vaccine also promoted differentiation of lymphoid lineage cells, including pre-B, B and NK cells, likely by enhancing the G1 to S transition, TGF-β signaling and Wnt signaling. We also observed that expression of the transcription factors Fos, Klf4 and Klf6 was significantly upregulated in pre-B, B and NK cells after G1-C vaccination, suggesting that these factors underlie changes in lymphoid lineage cells. Compared with the T-cell-directed vaccine BNT162b4, our new G1-C vaccine not only boosted T cell immune response but also enhanced the antibodies and B cell responses and promoted lymphoid lineage differentiation.

In summary, we identified a SARS-CoV-2 membrane epitope (LVGLMWLSYFIASFRLFARTRSMWSFNPETNIL) that can enhance adaptive immune responses and increase protection efficiency relative to that of the traditional RBD vaccine. This membrane region has been predicted to be presented on HLA class I and class II^42^. Our vaccine was also capable of regulating lymphoid-myeloid bias in BM. Due to the highly effective immune response induced by G1-C vaccine, it may be possible to reduce the dose of SARS-CoV-2 mRNA vaccine. Studies found that spike mRNA vaccine could induce postimmunization adverse effects, including stroke, thrombosis, myocarditis, herpes zoster reactivation, etc^75,76^. These adverse effects may relate to LNP or spike protein^76–79^. G1-C vaccine has smaller antigen compared to BNT162b4 or spike vaccine, which may provide safer strategy than using the large antigens. Altogether, our findings offer new insight into COVID-19 vaccine design and optimization.

This study has several limitations that highlight important avenues for future investigation. First, while we demonstrated that G1-C-induced antibodies exhibit superior neutralizing activity against multiple SARS-CoV-2 variants in vitro, our in vivo challenge experiments were limited to the D614G variant due to resource constraints. Nevertheless, the robust cross-variant neutralization observed in vitro, combined with enhanced protection against D614G challenge, provides strong preliminary evidence for the vaccine’s broad protective potential.

Second, our experiments focused on a single mouse strain (C57BL/6) and discrete post-vaccination timepoints. While this approach enabled rigorous proof-of-concept evaluation, extended longitudinal studies across multiple strains and relevant animal models will be essential to fully define the durability and generalizability of G1-C-mediated immune enhancement. Importantly, the magnitude and consistency of effects observed at our evaluated timepoints—including 8.2-fold increases in RBD-specific antibodies and significant modulation of bone marrow hematopoiesis—suggest biologically meaningful and reproducible phenomena worthy of further characterization.

Third, our immunological analyses emphasized spleen and bone marrow rather than draining lymph nodes (dLN), which serve as primary sites for germinal center reactions following intramuscular vaccination. While splenic analyses provide valuable insights into systemic immune activation, future studies incorporating detailed profiling of Tfh cells, germinal center B cells, memory T cells, and plasma cells in dLN will be critical to understand how G1-C modulates early lymph node responses and how these local events contribute to the enhanced systemic immunity we observed. This represents a logical and high-priority extension of the current work.

Finally, our transcription factor target gene analysis relied on ChIP-Atlas 3.0 databases, which may provide incomplete coverage depending on cell type and experimental conditions. Direct ChIP-seq or CUT&RUN experiments in vaccine-stimulated bone marrow cells would provide more definitive mapping of Fos, Klf4, and Klf6 regulatory networks.

Importantly, this study establishes associations between G1-C vaccination and enhanced immune responses but does not definitively prove causal mechanistic links with trained immunity. However, we have provided substantial mechanistic insights that extend well beyond simple correlative observations: (1) identification of a novel immunogenic epitope that induces >8-fold enhancement of RBD-specific antibodies, (2) demonstration that G1-C vaccination modulates bone marrow hematopoiesis and lymphoid lineage development, (3) identification of activated TGF-β and Wnt signaling pathways through transcriptomic analysis, and (4) functional validation showing that pharmacological inhibition of these pathways significantly reduces lymphoid cell proportions.

Most significantly, while the vast majority of SARS-CoV-2 vaccine studies have focused exclusively on antigen-specific responses in peripheral lymphoid organs, our work reveals that mRNA vaccination also modulates the bone marrow immune compartment—a previously underappreciated dimension of vaccine biology. This discovery opens an entirely new avenue for understanding how mRNA vaccines shape systemic immunity and provides a conceptual framework for next-generation vaccine design that deliberately targets bone marrow immune remodeling.

We view this work as foundational proof-of-concept that identifies a novel immunological phenomenon and establishes a platform for detailed mechanistic investigation. The consistency and magnitude of our findings—combined with functional pathway validation—provide compelling evidence that G1-C-mediated immune enhancement operates through design-driven mechanisms beyond simple antigen expression. Future studies employing epigenetic profiling, chromatin accessibility analyses, and longitudinal functional validation will build upon this foundation to fully elucidate the mechanisms underlying trained immunity in the context of mRNA vaccination. By revealing this new layer of vaccine-induced immunity, our work makes a significant contribution that warrants publication and will guide future research in the field.

## METHODS

### Mice

All studies were approved by Institutional Review Board (IRB) protocols and performed in accordance with Institutional Animal Care and Use Committee (IACUC) guidelines at the University of California, San Diego. Male C57BL/6J (Jackson Laboratory) and K18-hACE2 (Jackson Laboratory) mice were used at 6–8 weeks of age. C57BL/6J mice and uninfected K18-hACE2 were housed under barrier and specific-pathogen-free (SPF) conditions in individually ventilated cages (Innovative individually ventilated cages, kept under positive pressure during the study) with a maximum of four animals per cage. Temperature and relative humidity in the cages and animal unit were kept at 20–24°C and 45–55%, respectively, and the air change (AC) rate in cages was 75 AC/hr. All materials were autoclaved prior to use. Infected K18-hACE2 mice were housed in La Jolla Institute for Immunology animal biosafety level 3 (ABSL3) facility.

### Cell culture

Vero (ATCC, CCL-81) and HEK293T (ATCC CRL-3216) cell lines were maintained in Dulbecco’s modified Eagle’s medium (DMEM, Gibco) supplemented with 10% fetal bovine serum (FBS, Gibco) and penicillin (100 U/ml)- streptomycin (100 mg/mL) (Thermo Fisher Scientific). Lines were tested and confirmed to be mycoplasma-negative.

### Western blot

HEK293T cells were transfected with 0.5 μg of each mRNA. After 24 hrs, cells were lysed in RIPA buffer containing protease and phosphatase inhibitor cocktails (Thermo Scientific, 78441) and centrifuged at 12,000 × g for 10 min at 4 °C. All samples were boiled 10 min in loading buffer before being separated on 4-12% NuPAGE Bis-Tris gels and transferred to PVDF membranes. Membranes were blocked with 5% nonfat dry milk, incubated with primary antibodies at 4 °C overnight and then incubated 1 hr with horseradish peroxidase-conjugated goat anti-rabbit IgG (Cell Signaling Technology, 7074) at RT. Blots were visualized using ECL substrate (Thermo Scientific, 35050). Primary antibodies used were: anti-GAPDH (Proteintech, HRP-60004), anti-SARS-CoV-2 Spike RBD (Sino Biological, #40592-T62)

### Vaccine design

To design the vaccine, we incorporated a non-spike sequence immediately downstream of the RBD (ancestral virus) sequence within the SARS-CoV-2 spike protein. A linker sequence (GGGGS x 3) was introduced to bridge the RBD and newly inserted sequence. That linker was selected to preserve native folding of both the front and back regions of the protein, ensuring minimal interference with the immune response to the antigen.

### Immunization and endpoint collection

Male C57BL/6J mice (aged 6-8 weeks) were purchased from the Jackson Laboratory and housed based on regulatory standards of the University of California, San Diego. Mice were randomly allocated to experimental groups. For immunization, the mRNA-LNP vaccine was administered via intramuscular (IM) injection into the quadriceps muscle. Each 50 μl injection was delivered using a 28G needle insulin syringe. Mice were primed by IM injection of mRNA-LNP (10 μg mRNA) and boosted 2 weeks later with the same mRNA-LNP dose and administration route. Both vaccine doses were administered on the same side.

At specified endpoints, mice underwent humane euthanasia, initially via CO2 exposure, followed by a secondary euthanasia involving cervical dislocation. Cardiac puncture was then conducted to obtain whole blood, which was transferred into vacutainer Blood Collection Tubes containing K2 EDTA. Peripheral blood mononuclear cells (PBMCs) and plasma were isolated using SepMate™-15 (STEM CELL) tubes via density gradient centrifugation.

### Peripheral blood mononuclear cells (PBMCs) isolation

PBMCs were isolated from whole blood using SepMate™-15 (STEM CELL) tubes according to the manufacturer’s instructions with minor modifications. Briefly, 4.5 mL of density gradient medium was added to each SepMate™ tube. Blood samples were diluted 1:1 with PBS containing 2% fetal bovine serum (FBS) and gently mixed. The diluted samples were carefully layered onto the density gradient by pipetting down the side of the tube. Samples were centrifuged at 1200 × g for 10 min at RT with the brake on. Plasma was transferred to a new tube up to the white cell layer, and the PBMC layer was collected into a fresh tube. Collected PBMCs were washed twice with PBS + 2% FBS. Red blood cells were lysed by incubating cells in RBC lysis buffer (eBioscience) for 5 min at RT, and the reaction was stopped by adding ten volumes of PBS. Following a final wash, PBMCs were resuspended in complete RPMI medium (10% FBS, 0.5% penicillin/streptomycin).

### mRNA-LNP generation

All vaccines consist of 5′ and 3′ untranslated regions of human hemoglobin subunit alpha 1, IgE signal peptide, RBD coding sequences and non-spike epitope sequences. Sequences were codon-optimized and cloned into pBlueScript. DNA vectors were linearized, and mRNA synthesized in vitro using T7 polymerase (Cellscript, C-ASF3507), with UTP substituted with m1Ψ-5’-triphosphate (TriLink, N-1081). The 5’-Cap was added using a ScriptCap™ Cap 1 Capping System (Cellscript, C-SCCS1710). The poly(A) tail was added using a Poly(A) Tailing Kit (Cellscript, C-PAP5104H). mRNA was purified using LiCl (Sigma, SLCC8730, 2M final concentration).

To generate mRNA-LNPs, (6Z,9Z,28Z,31Z)-heptatriaconta-6,9,28,31-tetraen-19-yl 4-(dimethylamino)butanoate (D-Lin-MC3-DMA), 1, 2-distearoyl-sn-glycero-3-phosphocholine, cholesterol, and 1,2-dimyristoyl-rac-glycero-3-methoxypolyethylene glycol-2000 (DMG-PEG2000) were combined in ethanol at a molar ratio of 50:10:38.5:1.5. LNPs were formed by a self-assembly process in which the lipid mixture was rapidly mixed with the relevant indicated mRNA in 100 mM sodium acetate (pH 4) and incubated at 37°C for 15 min. mRNA-LNPs were then dialyzed in PBS (PH 7.4) at 4°C overnight. mRNA concentration was assessed using a RiboGreen RNA Assay Kit (Thermo Fisher Scientific, R11490). Particle size was analyzed using a dynamic light scattering apparatus (Malvern NANO-ZS90 Zetasizer).

### Preparation of splenocytes and bone marrow stem cells

For splenocytes, single-cell suspensions were prepared from spleen by homogenizing them through 70 μm cell mesh using a syringe plunger. Splenocytes were washed with excess DPBS and pelleted by centrifugation at 500 g for 5 min at RT. Supernatants were discarded.

For stem cells, BM was isolated from tibia and femur bones following removal of surrounding muscle. Joints were cut using a scalpel and the exposed BM was flushed from the ends of bones using a 25-gauge needle and a 3 ml syringe filled with 2% FBS PBS. Clumps were gently disaggregated using a needle-less syringe and passed through a 70 μm cell strainer. The cell suspension was pelleted by centrifugation at 500 g for 5 min at room temperature.

Erythrocytes from splenocytes and BM were then lysed 5 min in RBC lysis buffer (eBioscience) at RT. The reaction was stopped by adding 10 volumes of media containing 2% FBS. After another wash, cells were resuspended in complete RPMI medium (10% FBS, 0.5% penicillin/streptomycin) and passed again through 70 μm cell mesh. Cells were counted, and after resuspension in 1 mL of Bambanker Serum Free Cell Freezing Medium, frozen in liquid nitrogen.

### Flow cytometry

To analyze T cell activation in either PBMCs and splenocytes, 0.5 × 10^6^ cells were transferred to 96-well v-bottom cell culture plates and stabilized 4 hrs in a 37°C humidified incubator with 5% CO2. Cells were then stimulated with PepTivator® SARS-CoV-2 Prot_S1 (Miltenyi Biotec, 130-127-041) and PepTivator® SARS-CoV-2 Prot_M (Miltenyi Biotec,130-126-703) at the concentration as described in instructions for 16 hrs in the presence of brefeldin A (5.0 μg/mL).

To detect RBD specific B cells, we used Recombinant SARS-CoV-2 Spike RBD Alexa Fluor® 488 Protein (R&D systems, AFG10500) and APC/Cyanine7 conjugated SARS-CoV-2 Spike RBD. The Biotinylated Recombinant SARS-CoV-2 S Protein RBD (Biolegend, 793904) was incubated with APC/Cyanine7 Streptavidin (Biolegend, 405208) at a 3:1 ratio for 1 h at 4 °C^53^.

Cells were washed twice in PBS and then processed for flow cytometry staining. Specifically, cells were stained with fixable LIVE/DEAD Aqua to exclude dead cells and blocked with CD16/32 monoclonal antibody (Thermo Fisher Scientific, catalog# 14-0161-81). Cells were stained with different combinations of Live/Dead Aqua and surface markers antibody. Cells were permeabilized for intracellular marker staining. The antibodies used in staining are shown in supplementary Table 1. Stained samples were measured or sorted using a BD FACSCanto RUO-GREEN system. Data was analyzed using FlowJo v10 software.

### Isolation and sorting for 10x scRNAseq

1 × 10^6^ BM cells were transferred to 96-well v-bottom cell culture plates for staining with fixable LIVE/DEAD Aqua, based on product instructions. Stained samples were sorted using a BD ARIA II sorter. For 10X, single cells were processed with chromium next GEM single cell 3’ HT reagent kits v3.1 (dual index) (10X Genomics, Pleasanton, CA) following the manufacturer’s protocol. Approximately 33,300 cells were loaded to each channel with an average recovery rate of 20,000 cells. Libraries were sequenced on NovaSeq S4 (Illumina, San Diego, CA).

### Enzyme-linked immunosorbent assay (ELISA)

Total IgA, IgM and IgG were analyzed using a Mouse ELISA kit (Invitrogen). Pre-coated plates were incubated with 2-fold serial dilutions of standard and 1000-fold dilutions of plasma samples for 2 hrs at room temperature on a shaker and then washed four times. Next, plates were incubated with HRP-conjugated detection antibody for 2 hrs at room temperature on a shaker, washed four times, and incubated with substrate solution. The reaction was stopped by adding stop solution after 15 mins incubation. Plates were read at OD450. Standard curves were constructed using standard OD values for concentration analysis.

Anti-RBD IgA and IgM were analyzed using a Mouse anti-SARS-CoV-2 Spike RBD ELISA kit (Creative Diagnostics). Anti-RBD IgG was analyzed using a Mouse Anti-SARS-CoV-2 Antibody IgG Titer Serologic Assay Kit (ACROBiosystems). Assay procedures followed those for total immunoglobulin analysis.

### Pseudovirus production

HEK293T cells were transfected 24 hrs with plasmids expressing respective spike proteins from SARS-CoV-2 Ancestral (WT), or Beta, Delta or Omicron variants and then infected with VSV pseudovirus (MOI=5) for 1 hr. Infected cells were washed four times with medium and incubated for an additional 24 hrs. Culture supernatants were collected, centrifuged to remove cell debris, filtered using a 0.22-μm filter and stored as aliquots at −80°C.

### Pseudovirus neutralization assay

Neutralization activity of plasma was measured using a single-round pseudovirus infection of Vero cells. Cells were seeded in 96-well plates at 1 × 10^4^ cells/well and cultured overnight. Heat-inactivated plasma was serially diluted five-fold with DMEM, mixed with pseudovirus, and then incubated 1 hr at 37°C prior to addition to Vero cells for infection. The culture medium was refreshed after 12 hrs and incubated for an additional 48 hrs. Luciferase activity was measured using a Bright-Glo firefly luciferase kit (Promega). Each diluted sample was tested in duplicate wells. Neutralization activity was calculated using the luciferase luminescence value with the following formula: 100 × (1 – (value with mAb – value in “non-infected”)/(value in “no mAb” – value in “non-infected”)).

### Viral load measurement

Tissue viral loads were determined by quantitative PCR (qRT-PCR) of mouse lung, heart, kidney, brain and spleen tissues using DNA/RNA shield (Zymo Research) after virus challenge experiments. After homogenization, RNA was extracted using a Quick-DNA/RNA Miniprep Plus Kit (Zymo Research). Reverse transcription and qRT-PCR were performed using an iTaq Universal Probes One-Step Kit (Bio-Rad) on Roche LightCycler480 System. Viral loads were quantified using a SARS-CoV-2 Standard (Bio-Rad). CDC N1 forward and reverse primers, and probe were as follows:

Forward primer (N1): 5′- GACCCCAAAATCAGCGAAAT −3′;
Reverse primer (N1): 5′- TCTGGTTACTGCCAGTTGAATCTG −3′;
Probe: 5′-FAM-ACCCCGCATTACGTTTGGTGGACC -BHQ1-3′.

### Viral challenge in K18-hACE2 mice

The K18-hACE C57BL/6J mice (strain: 2B6.Cg-Tg(K18-ACE2) 2Prlmn/J) (5 male per group, aged 6-8 weeks) were used for virus challenge experiments. Animals were housed in an animal biosafety level 3 containment laboratory for viral infection and manipulation. Mice were intranasally inoculated with 1 × 10^4^ plaque-forming units (PFUs) of live SARS-CoV-2 D614G virus, (See comment in main figure 7 legend about this procedure.) and body weights and clinical scores were monitored 6 days. Serum was collected for antibody analysis. Lung, brain, heart, kidney and spleen tissues were harvested for viral load detection at 6 days after the challenge, and animals were euthanized with a low dose of isoflurane.

All animal experiments were approved by the Institutional Animal Care and Use Committee at the La Jolla Institute for Immunology (LJI) ABSL3 (protocol number AP00001242) and strictly conducted based on the National Institutes of Health Guide for the Care and Use of Laboratory Animals.

### Histopathology and Immunofluorescence

At necropsy, the left lung was collected and placed in 10% neutral buffered formalin for histopathologic analysis. Tissue embedding, sectioning, hematoxylin and eosin (H&E) staining, and slide scanning were completed at Moores Cancer Center, UC SanDiego.

### Single cell RNA-seq processing

Single cell RNA samples were processed using Cellranger (version 6.0.1) with the reference genome mm10. Individual samples were processed and analyzed in Seurat (version 4.3) for quality control. Cells with <500 or >6000 expressed genes or >15% mitochondrial genes were removed.

Doublets were detected and removed from each sample using Scrublet with an automatic threshold, considering an expected doublet rate of 6% for all samples. Ambient RNA contamination was accounted for by running SoupX on each sample using its automated algorithm to estimate contamination rates. Gene expression count matrices were adjusted for predicted contamination and rounded to integers (adjustCounts(sc, roundToInt=TRUE)). Adjusted counts were used for both clustering and downstream analysis.

### Reference mapping

After individual sample quality control, high-quality barcodes from scRNA-seq were merged and reference mapped using Seurat (version 4.3). The reference map selected was from the following publication^57^: doi: 10.1038/s41556-019-0439-6.

Gene expression of both reference and query samples was normalized (NormalizeData) and scaled (ScaleData). Neighbors were identified in the reference map PCA space (FindNeighbors), and TransferAnchors were used to project the query to the reference UMAP and transfer cell type labels from the reference to the query. Cells with a prediction score <0.5 for label transferring were removed.

### Differential gene expression analysis

To determine changes in gene expression, we performed differential analysis using DESeq2. Pseudo-bulk matrices for each cell type were derived by summing the counts for all barcodes of a sample for each gene on a per cell type basis. These matrices were created using our SoupX-corrected and rounded expression count matrix. Genes were only tested if present in at least 2 samples per test condition and if there was a total of at least 10 counts across all tested conditions. Multiple test corrections were performed using the Benjamini & Hochberg method, with a significance threshold of padj=0.1.

### Pathway analysis

To assess canonical pathway enrichment, we performed gene set enrichment analysis (GSEA). Using results from our differential expression analysis, input genes were ranked using the DESeq2 STAT column, and fGSEA was run using both Wikipathways and REACTOME databases [parameters: eps=0.0, minSize=10, maxSize=500]. Enriched pathways were filtered using an FDR cutoff of 10%.

### Volcano plots

Differentially expressed genes (DEGs) identified in cells from mice immunized with G1-C versus RBD were visualized in volcano plots and shown as significantly upregulated or downregulated, versus non-significant differences. ChIP-Atlas 3.0 was used to identify Fos, Klf4 and Klf6 target genes at TSS ±5k from M. musculus (mm10). We selected data from MEFs for Fos, BM cells for Klf4, and oligodendrocytes for Klf6, and target genes were filtered by MACS q-value threshold of 0.05. Transcription factor target genes and DEGs (p-adj < 0.05) were overlapped and illustrated on the volcano plot, with the top 20 DEGs labeled.

### Cell cycle analysis

Cell cycle dynamics were analyzed using FxCycle™ Far Red Stain. 0.5 × 10^6^ BM cells were stained with different combinations of Live/Dead Aqua and specific antibody combinations. After staining with markers of interest, we added 1 μl FxCycle™ Far Red stain (5 μM) and 1 μl of RNase A (100 mg/mL) to each flow cytometry sample and mixed well. Cells were then incubated for 30 minutes at 4°C, while protected from light. Samples were analyzed in a flow cytometer without washing. Data was analyzed using FlowJo v10 software.

### Real-time qPCR

Total cellular RNA was extracted by TRIzol reagent (Invitrogen) and then reverse transcribed using using an iScript cDNA Synthesis Kit (Bio-Rad). RT-PCR was performed using 2× SYBR Green mix (Bio-Rad) and run on LightCycler480. PCR cycling conditions were: 95 °C for 5 min followed by 45 cycles of 95 °C for 10 s, 60 °C for 10 s, and 72 °C for 10 s. Data were analyzed using a SYBR green-based system (Toyobo), and semi-quantified and normalized with glyceraldehyde-3-phosphate dehydrogenase (Gapdh). Gapdh, Fos, Klf4, and Klf6 forward and reverse primers used were as follows:

Gapdh-F 5′-TGACCTCAACTACATGGTCTACA-3’; Gapdh-R 5′-CTTCCCATTCTCGGCCTTG-3’;
Fos-F 5’-CGGGTTTCAACGCCGACTA-3’; Fos-R 5’-TGGCACTAGAGACGGACAGAT-3’;
Klf4-F 5’-ATCCTTTCCAACTCGCTAACCC-3’; Klf4-R 5’-CGGATCGGATAGCTGAAGCTG-3’;
Klf6-F 5’-GTTTCTGCTCGGACTCCTGAT-3’; Klf6-R 5’-TTCCTGGAAGATGCTACACATTG-3’;
Prkcq-F 5’-TCCGCCAGCATCCTTTGTTT-3’; Prkcq-R 5’-TCCTTGTCGAAATTGCTACAGTC-3’;
Prkcb-F 5’-GTGTCAAGTCTGCTGCTTTGT-3’; Prkcb-R 5’-GTAGGACTGGAGTACGTGTGG-3’;
Smad1-F 5’-GCTTCGTGAAGGGTTGGGG-3’; Smad1-R 5’-CGGATGAAATAGGATTGTGGGG-3’.

### Statistical analysis

Statistical analyses were performed with GraphPad Prism 10. ANOVA with Šidák’s correction was used for post-hoc testing, and multiple t-tests were analyzed using the two-stage linear step-up procedure of Benjamini, Krieger, and Yekutieli to control the false discovery rate^80^. Statistical significance was inferred if *P* < 0.05. *P* values are indicated by \**P* < 0.05; \*\**P* < 0.01; \*\*\**P* < 0.001; \*\*\*\**P* < 0.0001.

## Data and code availability

Single-cell RNA sequencing data are deposited on GEO: GSE281662.

The code used for data analysis is available on GitHub: https://github.com/jingw1072/mice-BM-scRNAseq-analysis-code.

## Acknowledgements

We thank Dr. Kristen Jepsen of the Institute of Genomic Medicine at UCSD for help with scRNA-seq, Dr. Neal Sekiya and Ms. Tara Rambled at the Center for AIDS Research at UCSD for flow cytometry analysis, and members of the Rana lab for helpful discussions and advice. This publication includes data generated at the UC San Diego IGM Genomics Center utilizing an Illumina NovaSeq 6000 purchased with funding from a National Institutes of Health SIG grant (#S10 OD026929). This work was supported in part by institutional funds and from grants from the National Institutes of Health (AI187411, AI125103, CA177322, CA030199, DA046171).

## Author contributions

JW designed and performed experiments, analyzed data, and wrote the manuscript draft; JM designed and performed experiments, and analyzed data; LT and SB analyzed the scRNA-seq data; AS, RM, JT, and LW performed experiments; DMS provided reagents, SS and KJG participated in experimental design, data analysis and interpretation; TMR conceived and planned the project and participated in experimental design, data analysis, data interpretation, and manuscript writing.

---

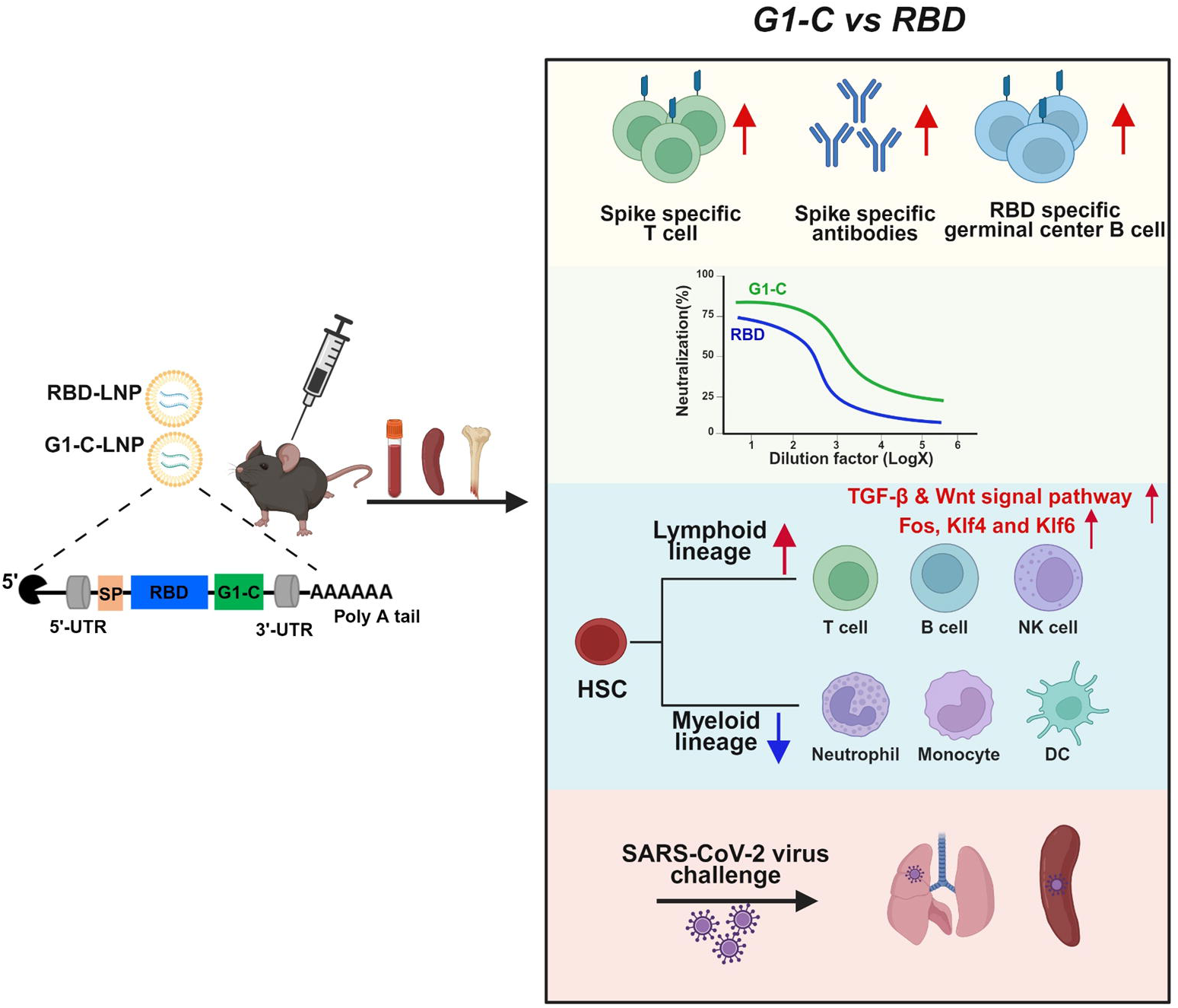

## Highlights

- After screening and in vivo immune analysis, we identified a membrane epoitope that significantly enhanced vaccine-induced immunity and protection in vivo.
- The new epitope, G1-C, vaccination modulates HSC to change the balance between lymphoid and myeloid lineage cells.
- G1-C mRNA vaccine promotes lymphoid lineage cells differentiation by upregulating TGF-β and Wnt signal pathways.
- G1-C mRNA vaccine increases levels of Band NK cells by upregulating Fas, K1f4, and K1f6 transcription factors.
- G1-C mRNA vaccine provides better protection efficiency against viral challenge in K18-hACE2 mice than that of the traditional RBD vaccine.

