## Supplementary Information for "A new mRNA antigen vaccine induces potent B and T cell responses and *in vivo* protection against SARS-CoV-2"

**Supplementary Fig 1**

A

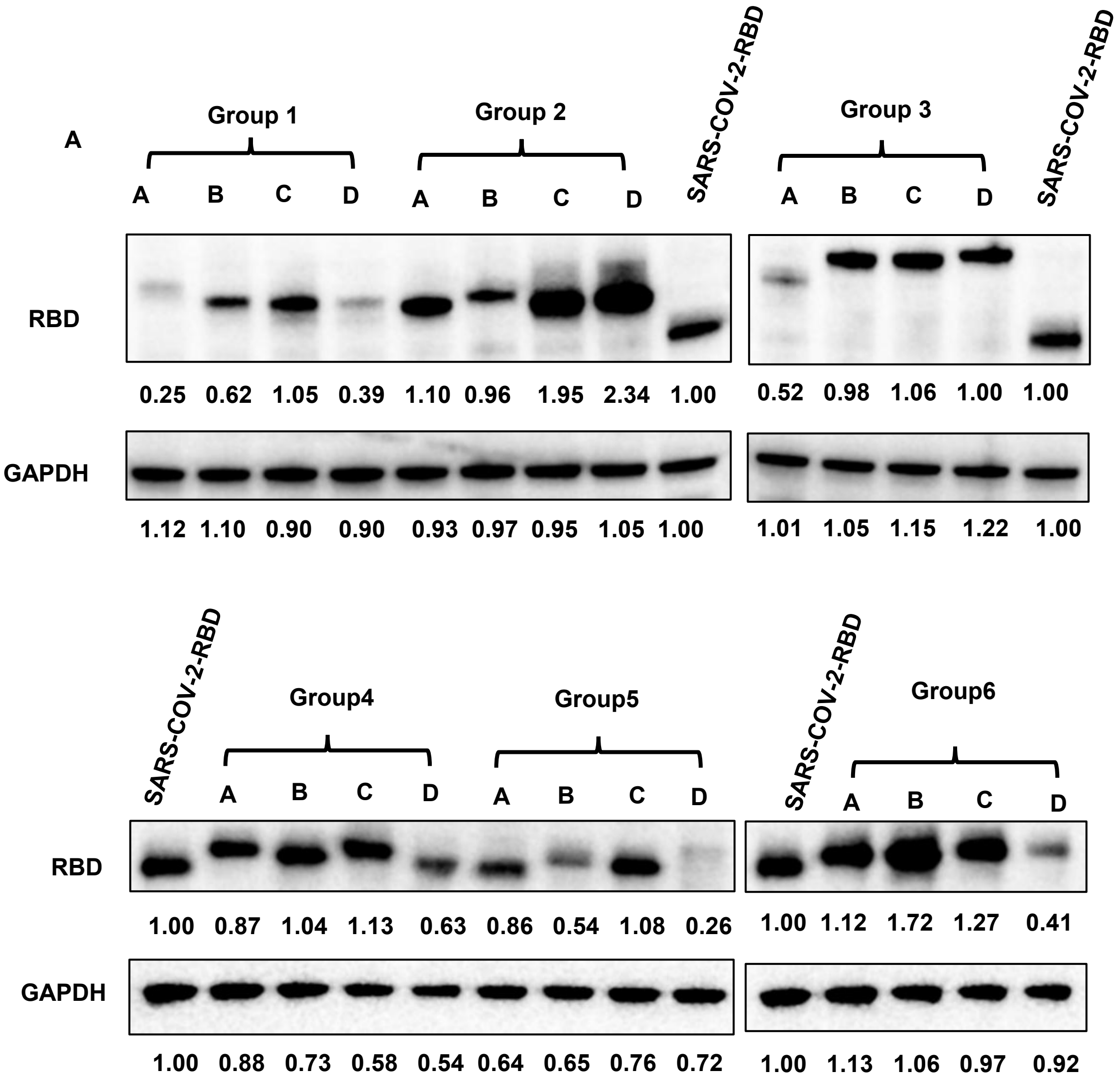


B

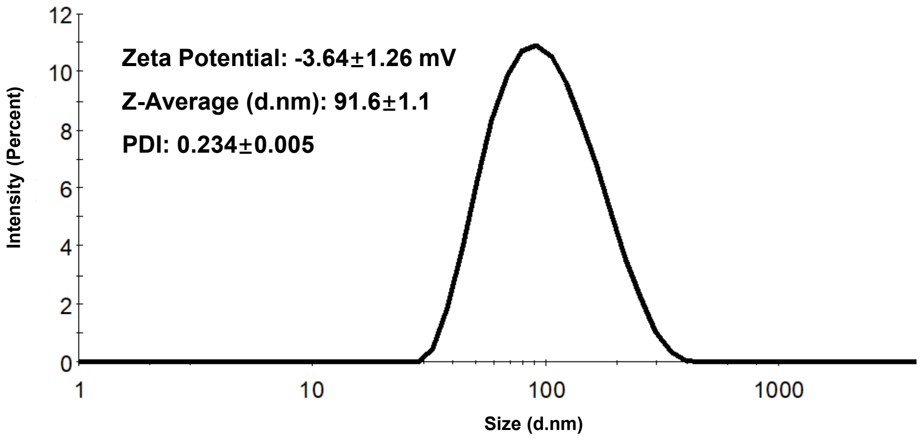

#### Supplementary Fig 2

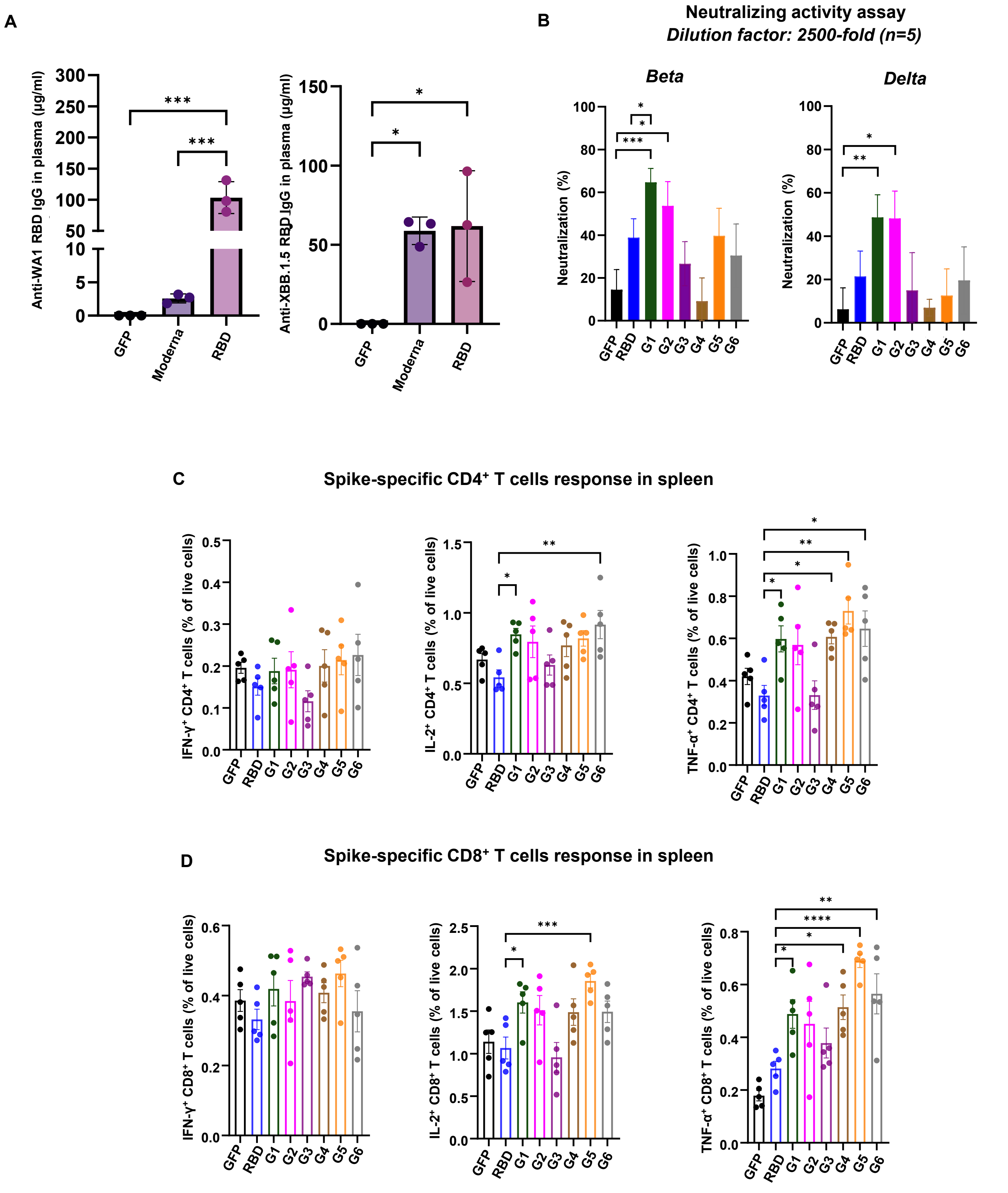

#### Supplementary Fig 3

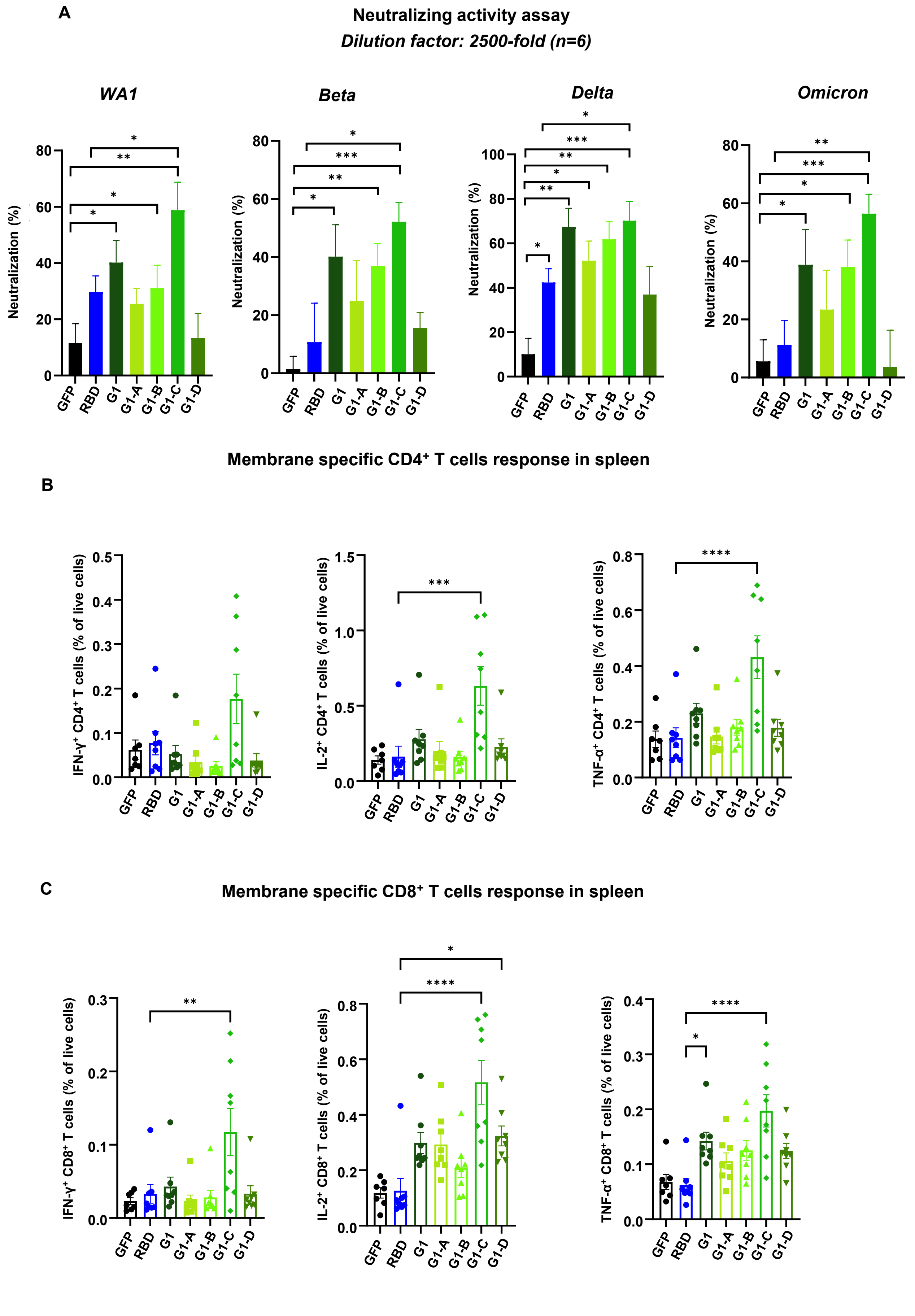

Supplementary Fig 4

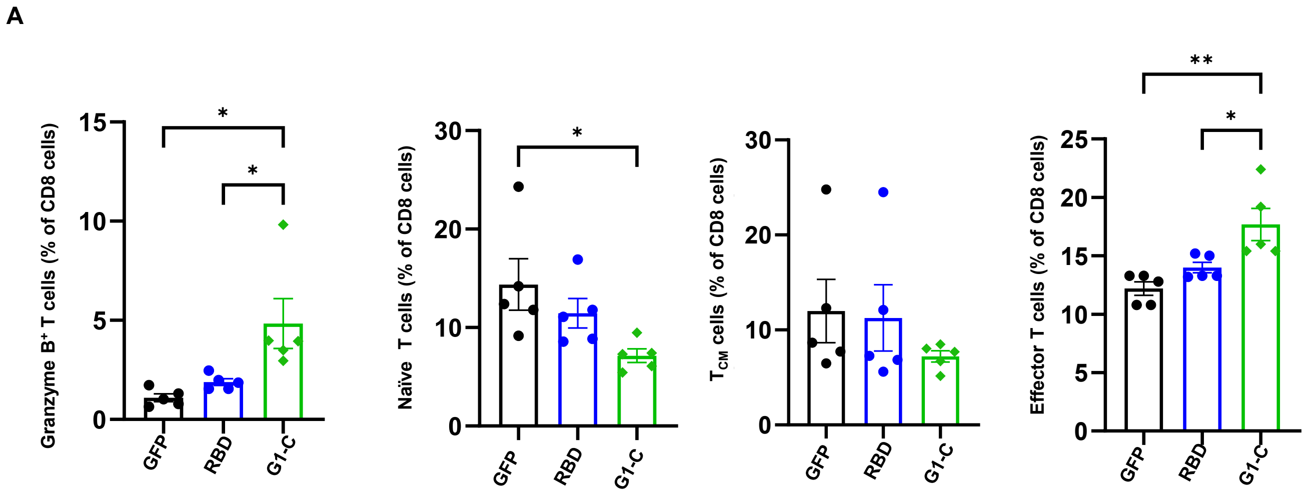


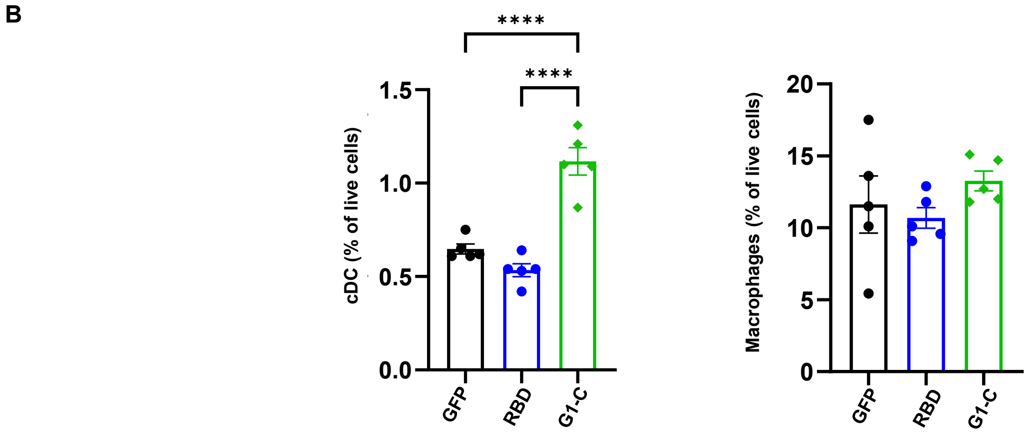

#### Supplementary Fig 5

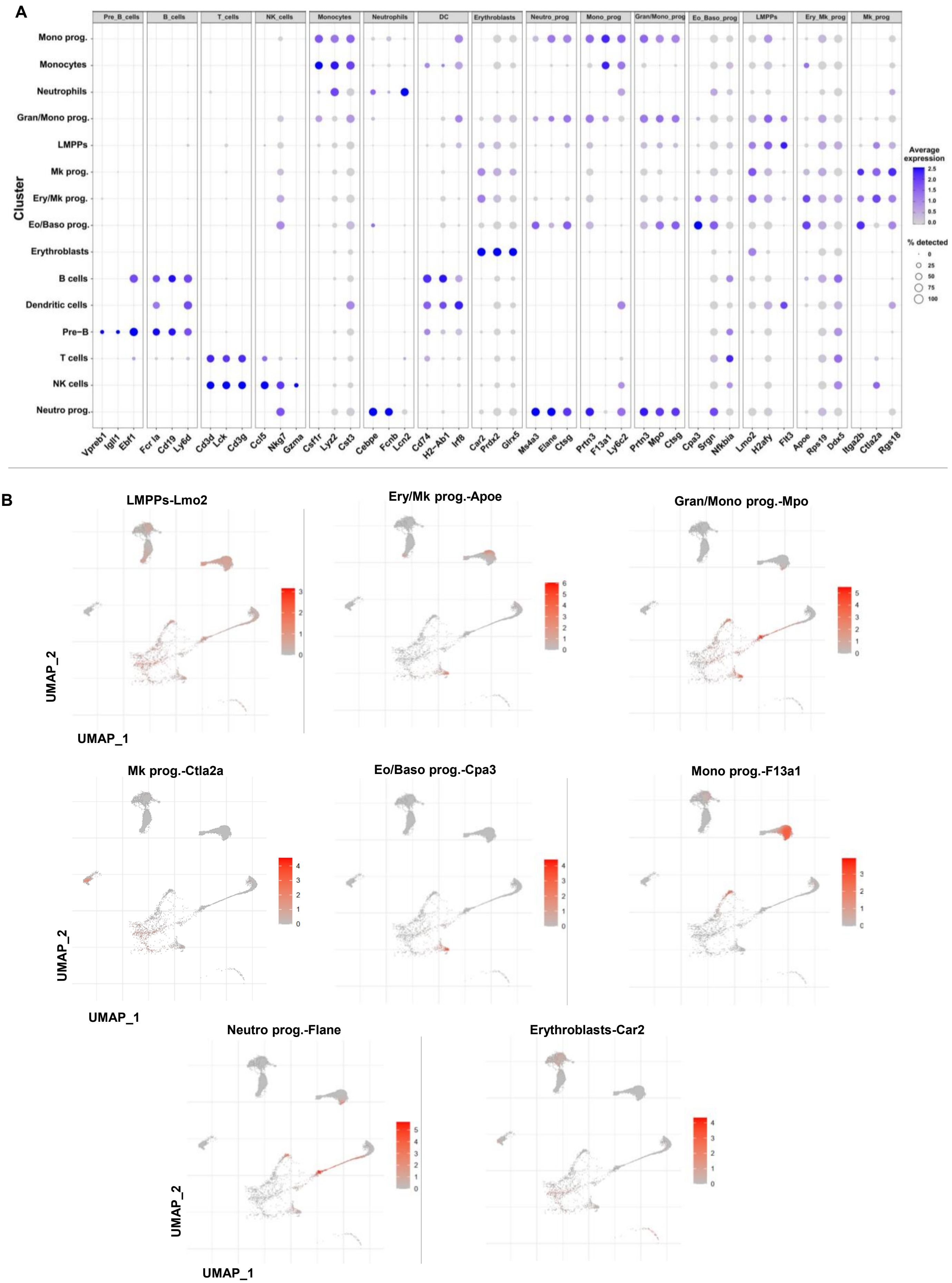

**Supplementary Fig 6**

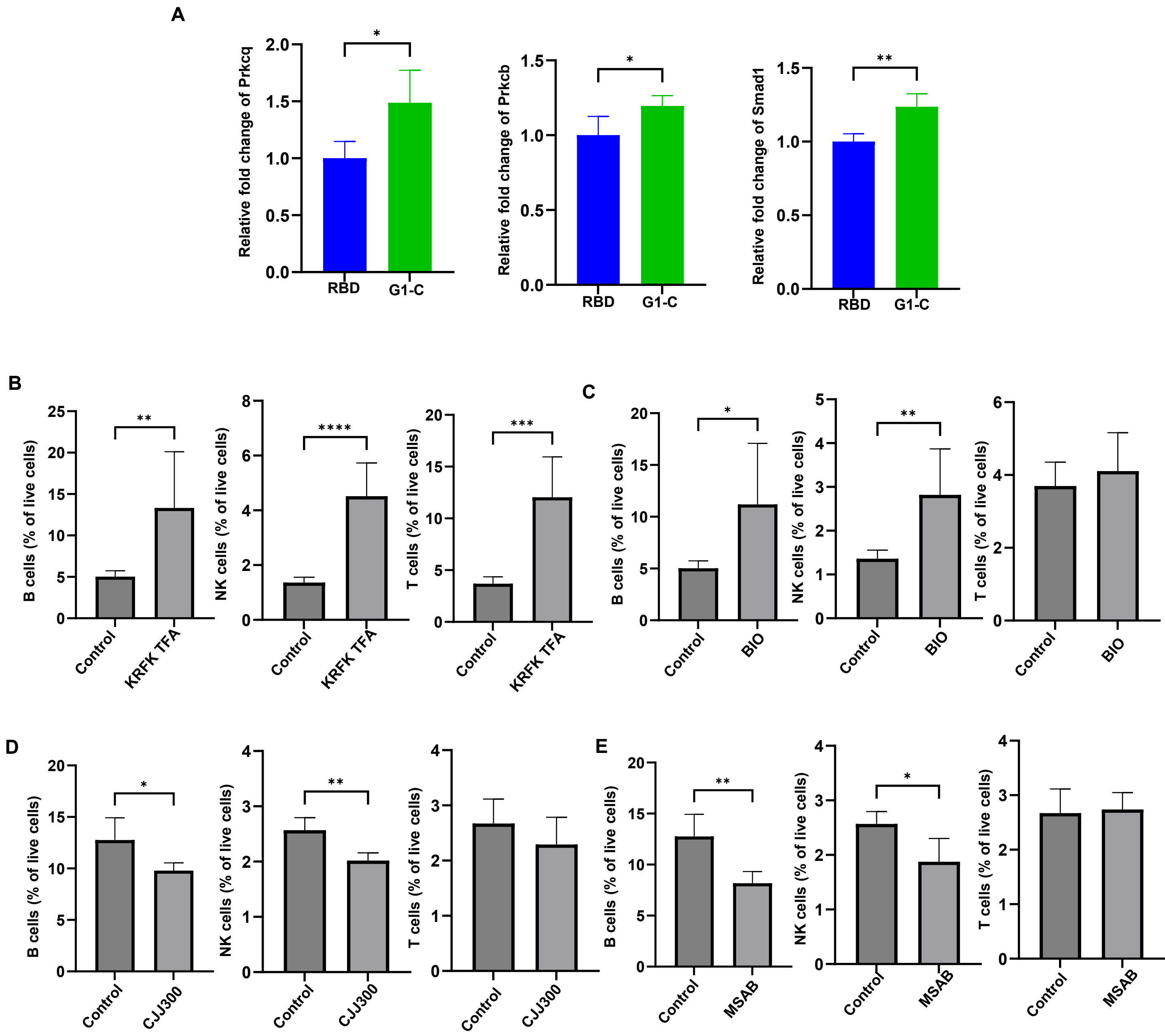

### Supplementary Fig 7

**A** **B cells**

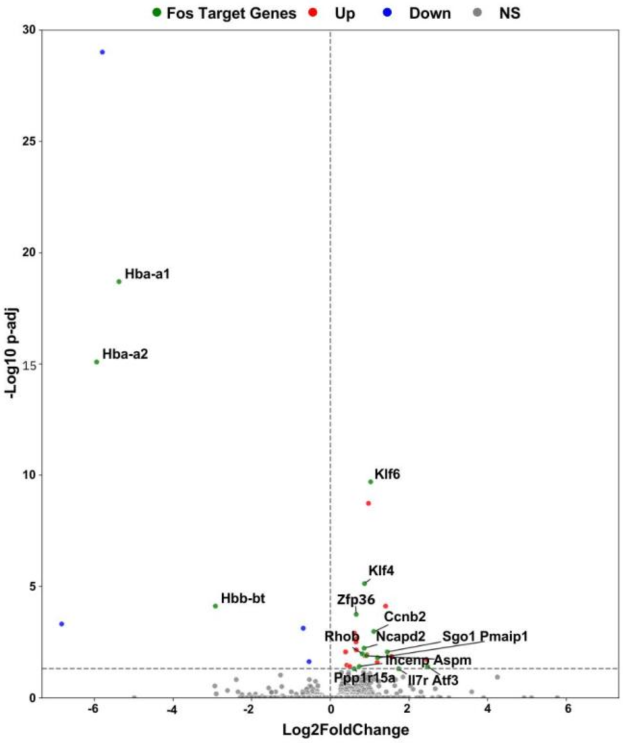


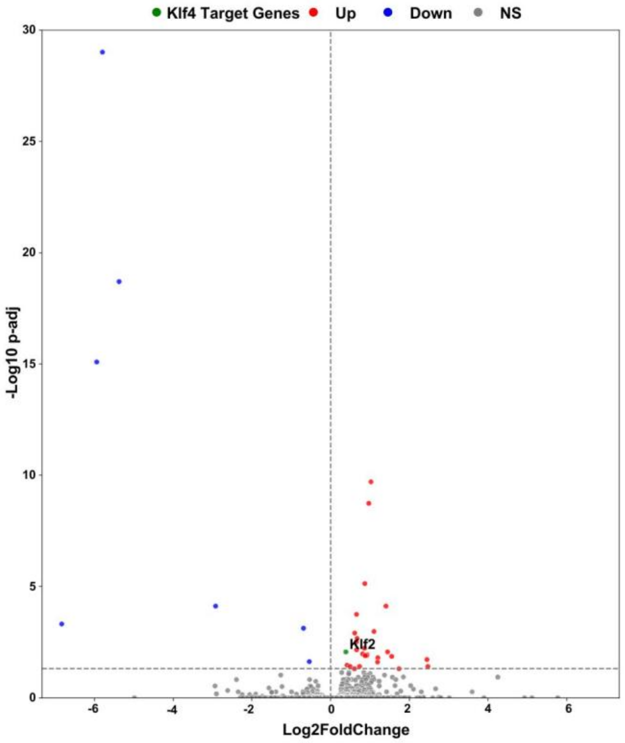


**B** **NK cells**

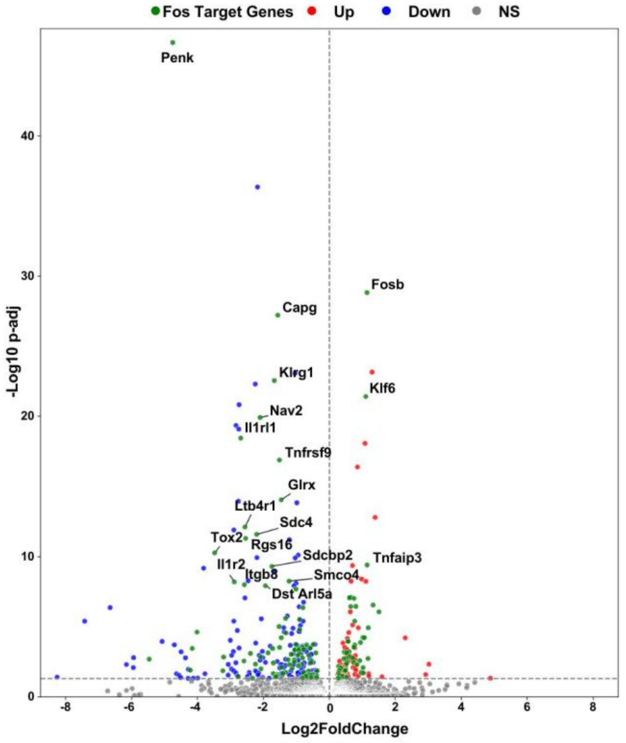


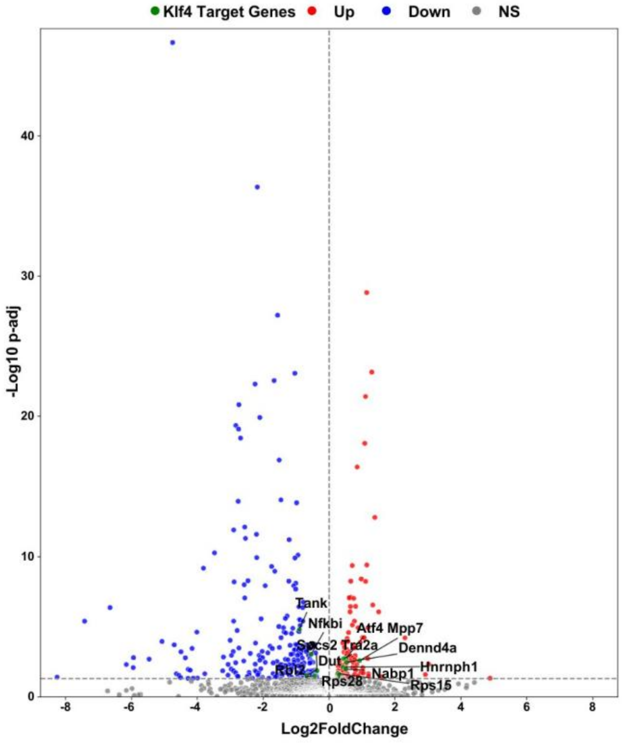

### Supplementary Fig 8

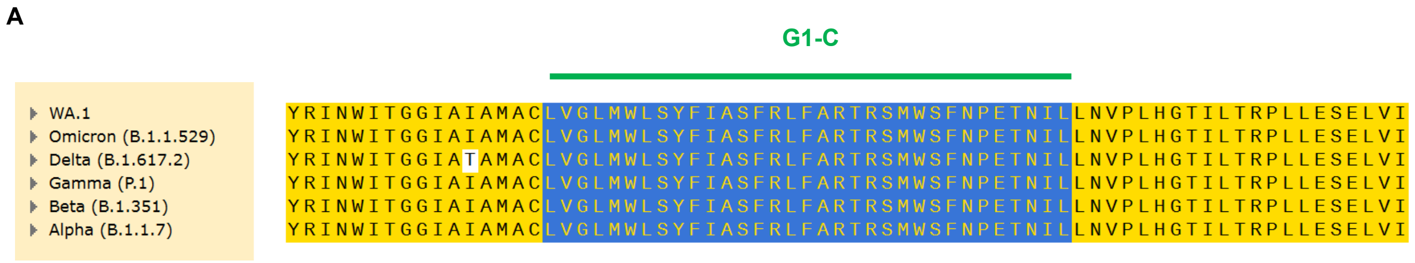

#### Supplementary Fig 9

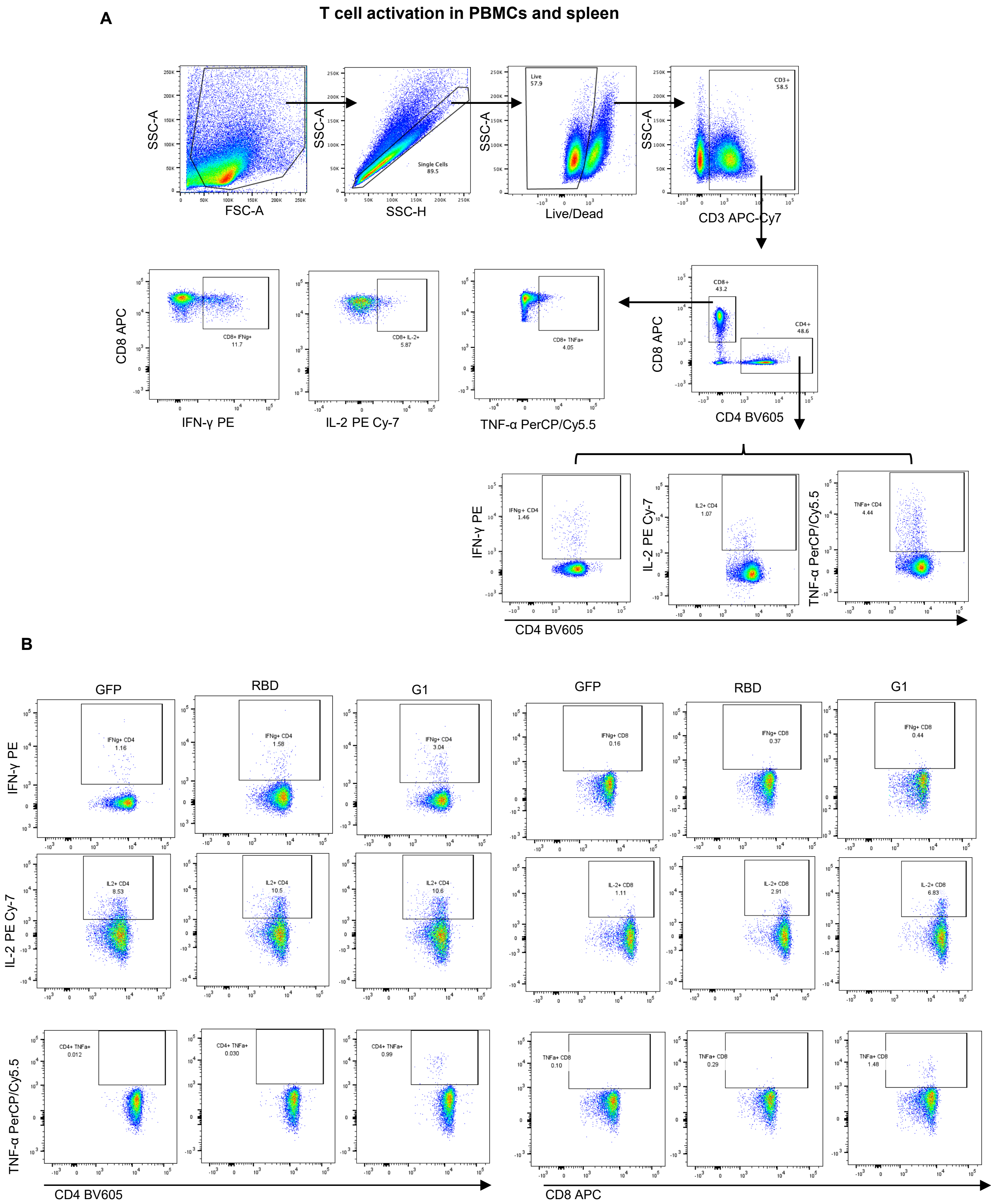

#### Supplementary Fig 10

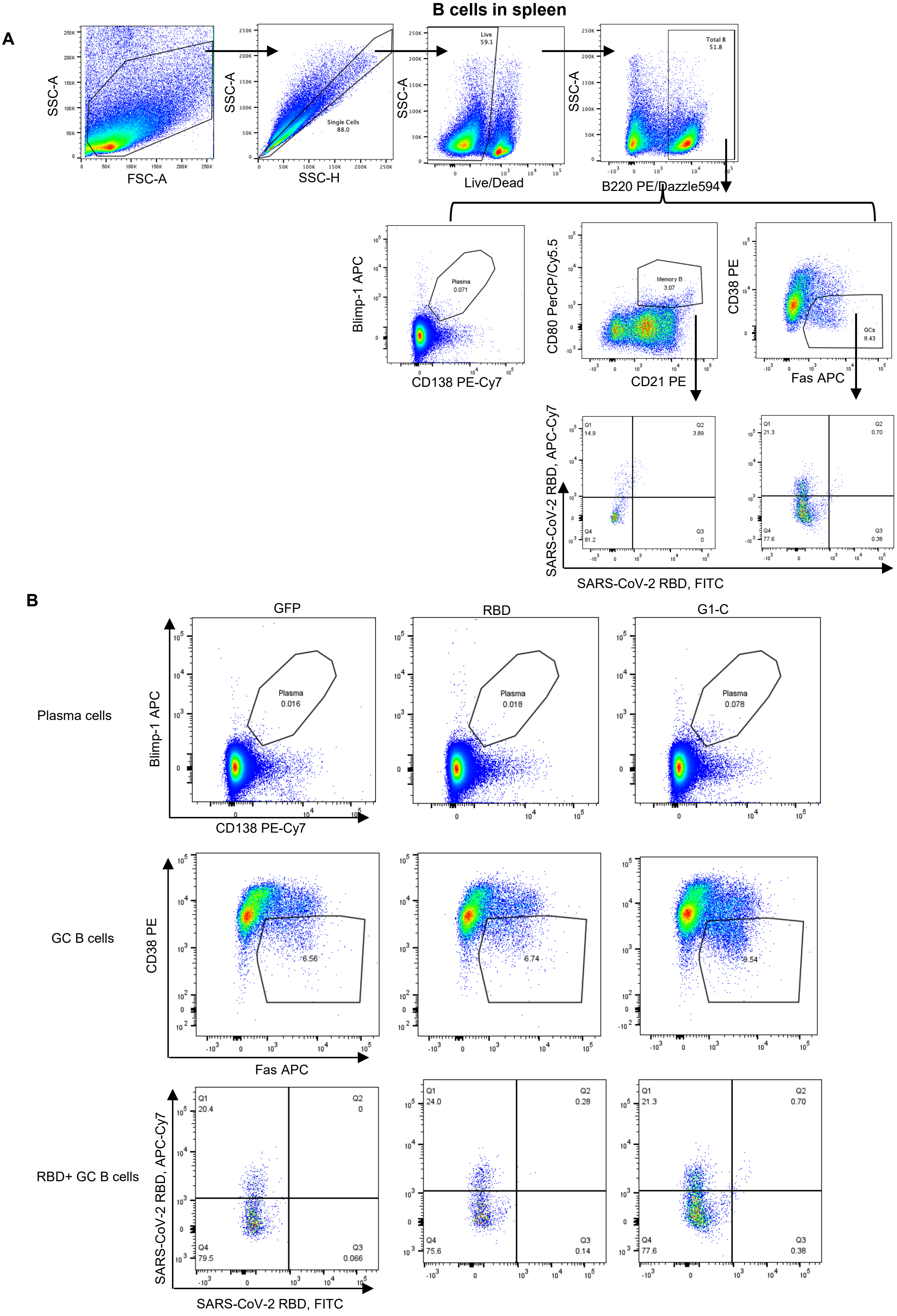

**Supplementary Fig 11**

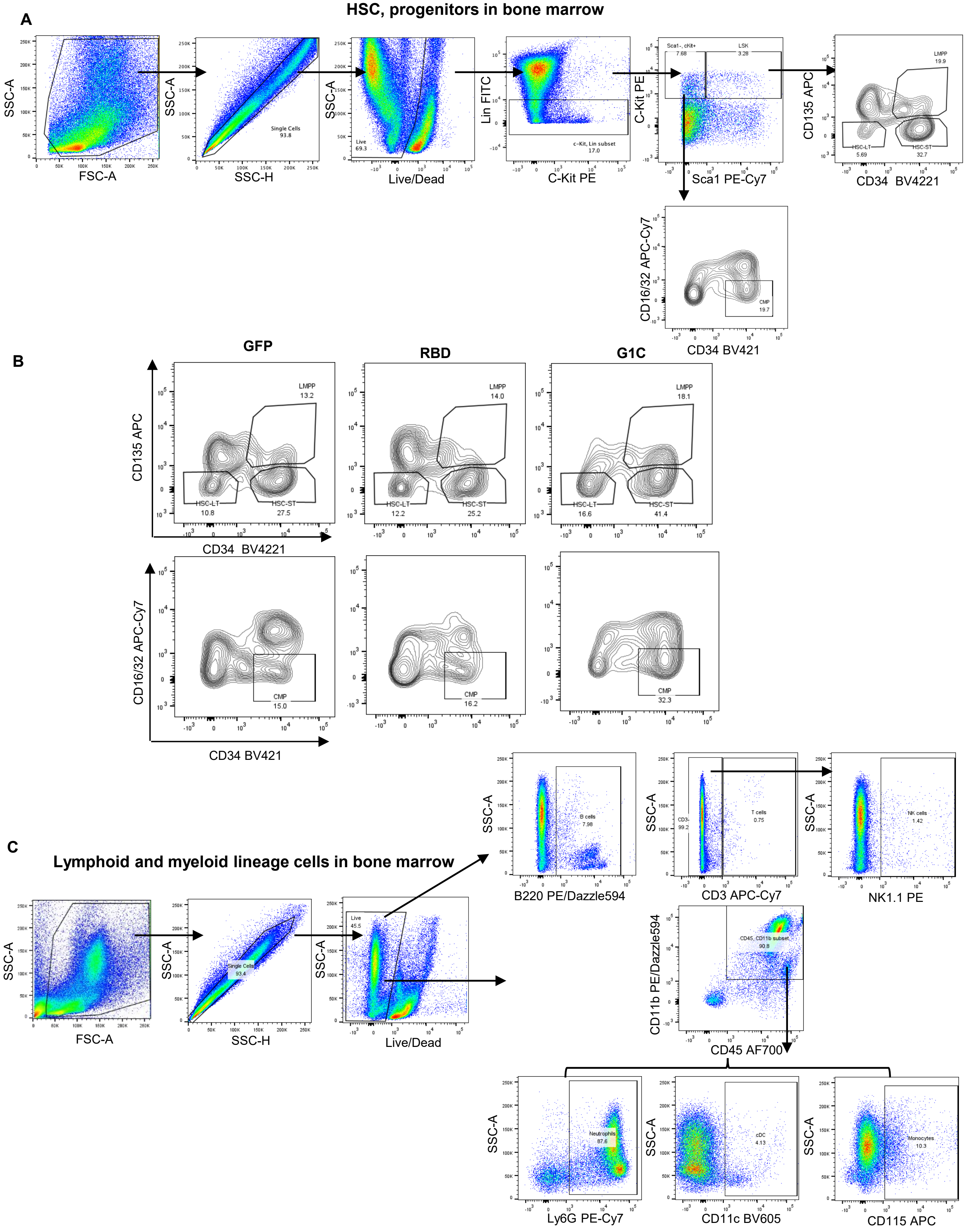

**Supplementary Fig 12**

**A** 

#### CD8 T cells in spleen

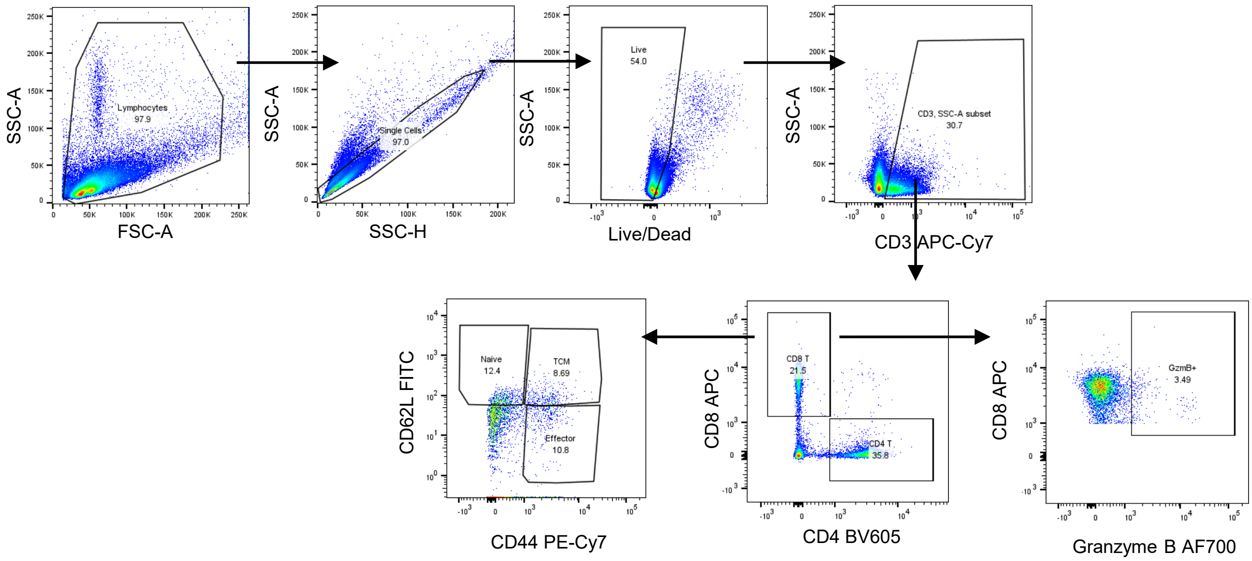


**B**

#### Macrophages and cDCs in spleen

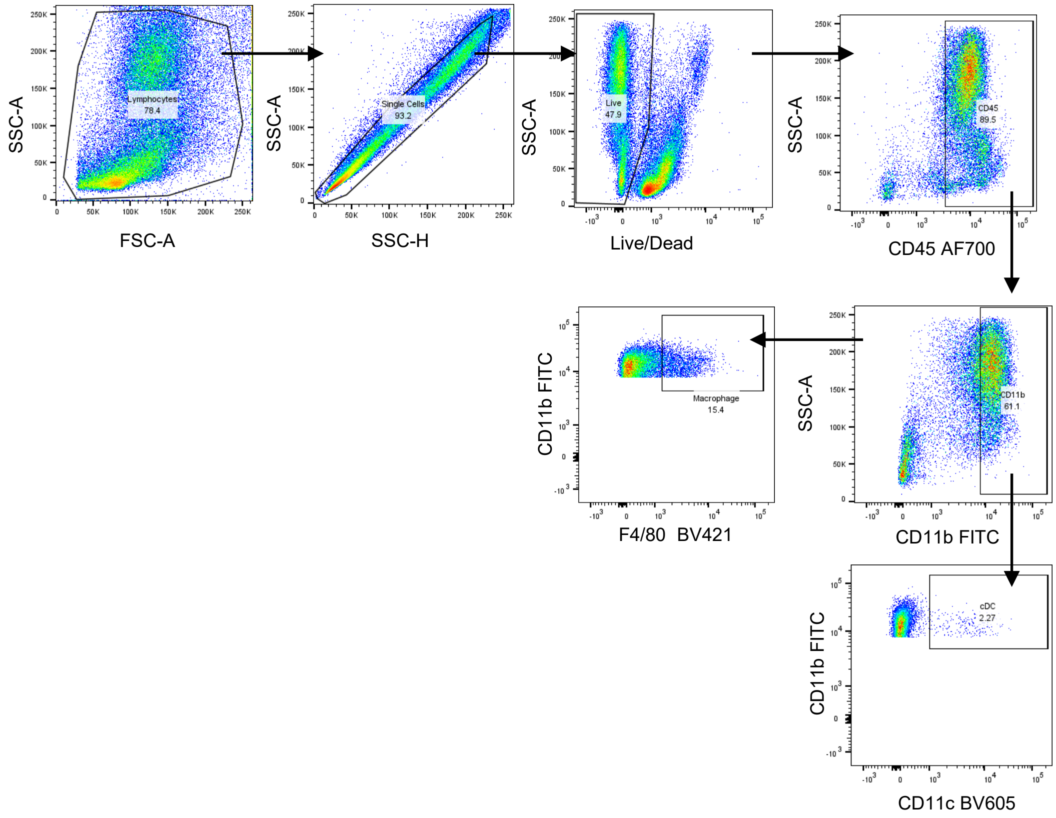

#### Supplementary Table 1

| Antibody | Source | Cat. No. |
|---|---|---|
| Brilliant Violet 421™ anti-mouse CD34 Antibody | Biolegend | 152208 |
| Brilliant Violet 605™ anti-mouse CD41 Antibody | Biolegend | 133921 |
| FITC anti-mouse Lineage Cocktail | Biolegend | 133302 |
| PE anti-mouse c-Kit Antibody | Biolegend | 105808 |
| PerCP/Cyanine5.5 anti-mouse CD150 Antibody | Biolegend | 115922 |
| PE/Cyanine7 anti-mouse Sca-1 Antibody | Biolegend | 122514 |
| APC anti-mouse CD135 Antibody | Biolegend | 135310 |
| APC/Cyanine7 anti-mouse CD16/32 Antibody | Biolegend | 101328 |
| Brilliant Violet 421™ anti-mouse F4/80 Antibody | Biolegend | 123137 |
| Brilliant Violet 605™ anti-mouse CD11c Antibody | Biolegend | 117334 |
| FITC anti-mouse CD3 Antibody | Biolegend | 100204 |
| FITC anti-mouse B220 Antibody | Biolegend | 103206 |
| FITC anti-mouse NK1.1 Antibody | Biolegend | 156508 |
| PE anti-mouse Siglec-F Antibody | Biolegend | 155506 |
| FITC anti-mouse/human CD11b Antibody, | Biolegned | 101206 |
| PE/Cyanine7 anti-mouse Ly-6G Antibody | Biolegend | 127618 |
| APC anti-mouse CD115 Antibody | Biolegend | 135510 |
| Alexa Fluor® 700 anti-mouse CD45 Antibody | Biolegend | 103128 |
| Brilliant Violet 605™ anti-mouse CD4 Antibody | Biolegend | 100451 |
| PE anti-mouse NK-1.1 Antibody | Biolegend | 108708 |
| PE/Dazzle™ 594 anti-mouse/human CD45R/B220 Antibody | Biolegend | 103258 |
| APC anti-mouse CD8a Antibody | Biolegend | 100712 |
| APC/Cyanine7 anti-mouse CD3ε Antibody | Biolegend | 100330 |
| Brilliant Violet 605™ anti-mouse CD4 Antibody | Biolegend | 100451 |
| IFN gamma Monoclonal Antibody (XMG1.2), PE | (eBioscience | 12-7311-82 |
| PerCP/Cyanine5.5 anti-mouse TNF-α Antibody | Biolegend | 506322 |
| PE/Cyanine7 anti-mouse IL-2 Antibody | Biolegend | 503832 |
| FITC anti-mouse CD21/CD35 Antibody | Biolegend | 123408 |
| PE anti-mouse CD38 Antibody | Biolegend | 102708 |
| PE/Cyanine7 anti-mouse CD138 (Syndecan-1) Antibody | Biolegend | 142514 |
| APC anti-mouse CD95 (Fas) Antibody | Biolegend | 152604 |
| Alexa Fluor® 700 anti-mouse CD23 Antibody | Biolegend | 101632 |
| PerCP/Cyanine5.5 anti-mouse CD80 Antibody | Biolegend | 104722 |
| Alexa Fluor® 700 anti-human/mouse Granzyme B Antibody | Biolegend | 396426 |
| FITC anti-mouse CD62L Antibody | Biolegend | 104406 |
| PE/Cyanine7 anti-mouse/human CD44 Antibody | Biolegend | 103030 |
| APC anti-mouse Blimp-1 Antibody | Biolegend | 150007 |

#### Supplementary Table 2

| Cell type | GFP1 | GFP2 | GFP3 | RBD1 | RBD2 | RBD3 | G1-C1 | G1-C2 | G1-C3 |
|---|---|---|---|---|---|---|---|---|---|
| B cells | 265 | 411 | 382 | 837 | 690 | 358 | 713 | 947 | 1345 |
| Dendritic cells | 688 | 456 | 460 | 1189 | 2236 | 1013 | 474 | 1161 | 1266 |
| Eo/Baso prog. | 18 | 23 | 27 | 34 | 24 | 12 | 96 | 80 | 33 |
| Ery/Mk prog. | 2 | 1 | 6 | 0 | 5 | 2 | 28 | 12 | 9 |
| Erythroblasts | 3 | 11 | 3 | 3 | 2 | 23 | 0 | 7 | 0 |
| Gran/Mono prog. | 7 | 10 | 12 | 17 | 5 | 8 | 68 | 90 | 48 |
| LMPPs | 44 | 40 | 63 | 56 | 54 | 43 | 220 | 200 | 128 |
| Mk prog. | 0 | 0 | 5 | 2 | 1 | 3 | 3 | 4 | 3 |
| Mono prog. | 6 | 11 | 6 | 16 | 9 | 3 | 77 | 89 | 49 |
| Monocytes | 1276 | 1902 | 1608 | 1823 | 2870 | 1367 | 3269 | 4982 | 2890 |
| Neutro prog. | 3 | 6 | 17 | 11 | 8 | 2 | 95 | 138 | 70 |
| Neutrophils | 7800 | 8501 | 11595 | 6465 | 5862 | 5125 | 6451 | 2351 | 3633 |
| NK cells | 236 | 229 | 471 | 201 | 314 | 282 | 583 | 1207 | 988 |
| pre-B | 297 | 1367 | 617 | 2401 | 1218 | 504 | 1710 | 3382 | 2310 |
| T cells | 14 | 12 | 25 | 19 | 16 | 25 | 27 | 24 | 34 |
| Total | 10659 | 12980 | 15297 | 13074 | 13314 | 8770 | 13814 | 14674 | 12806 |

**Supplementary Figure 1. Expression of mRNA constructs and characterization of mRNA-lipid nanoparticles (LNPs)**

(A) Western blot analysis of expression of all 24 mRNA constructs in HEK293T cells. Cells ($1\times10^5$) were transfected with 500 ng mRNA and collected for analysis 24 hrs later. Blots were probed with anti-SARS-CoV-2 Spike protein and anti-GAPDH. Grey intensity of the Western blotting strips was analyzed with software of Image J.

(B) Particle size and zeta potential of mRNA-LNPs, as determined by dynamic light scattering (DLS).

**Supplementary Figure 2. Analysis of neutralizing activity and memory T cell responses in spleen of mice immunized with G1 to G6 vaccines**

(A) XBB.1.5-RBD specific IgG titers in plasma, as quantified by ELISA.

(B) Neutralizing activity of plasma from mice immunized with indicated vaccines against Beta and Delta SARS-CoV-2 pseudoviruses. Mouse plasma was diluted 2500-fold for the assay (n=5).

(C, D) Spike-specific $CD4^+$ (B) and $CD8^+$ (C) T cell responses of mice immunized with indicated pooled vaccines, as measured in spleen and based on intracellular IFN-$\gamma^+$, IL-$2^+$ or TNF$\alpha^+$ staining. Asterisks represent statistical significance (*$p < 0.05$; **$p < 0.01$; ***$p < 0.001$; ****$p < 0.0001$) as determined by unpaired t-test (A) and one-way ANOVA (B, C). Error bars indicate standard error of the mean (SEM).

**Supplementary Figure 3. Immunization with G1-C mRNA vaccine promotes potent neutralizing activity and induces membrane-specific memory T cell responses in spleen**

(A) Neutralizing activity in plasma of mice immunized with indicated vaccines against WA1, Beta, Delta and Omicron SARS-CoV-2 pseudoviruses. Plasma was diluted 2500-fold for the assay.

(B, C) Membrane-specific $CD4^+$ (B) and $CD8^+$ (C) T cell responses seen following immunization with indicated vaccines, measured in spleen by intracellular staining of IFN-$\gamma^+$, IL-$2^+$ and TNF$\alpha^+$.

Asterisks represent statistical significance (*p < 0.05; **p < 0.01; ***p < 0.001; ****p < 0.0001) as determined by unpaired t-test (A) and one-way ANOVA (B, C). Error bars indicate standard error of the mean (SEM).

**Supplementary Figure 4. Immunization with G1-C mRNA vaccine promotes CD8 T cell and Classical Dendritic Cells (cDCs) responses in spleen**

(A) Analysis of the proportion of cytotoxic, naïve, central memory and effector CD8 T cells in spleen of mice immunized with indicated vaccines, based on flow cytometry (n=5).

(B) Analysis of the proportion of antigen-presenting cells including macrophages and cDCs in spleen of mice immunized with indicated vaccines, based on flow cytometry (n=5).

Asterisks represent statistical significance (*p < 0.05; **p < 0.01; ***p < 0.001; ****p < 0.0001) as determined by one-way ANOVA. Error bars indicate standard error of the mean (SEM).

**Supplementary Figure 5. Identification of bone marrow cell types by scRNAseq**

(A) The expression levels of 3 cell markers for each cell type are visualized in dot plots, related to Fig 4C.

(B) Expression of markers of progenitor cells highlighted on UMAP, related to Fig 4E.

**Supplementary Figure 6. TGF-β and Wnt signaling pathways regulate B and NK cell differentiation in bone marrow**

(A) RT-PCR analysis of expression of *Smad1*, *Prkcb* and *Prkcq* transcripts in BM cells ($5\times10^6$) of mice immunized with indicated vaccines. Values are normalized to Gapdh; shown is the relative fold-change (n=4).

(B) The proportion of living B, NK and T cells seen after treatment (48 hrs) of fresh isolated mouse BM cells ($2\times10^6$) with KRFK TFA (100 μM), a TGF-β agonist. Cells were analyzed by flow cytometry (n=7).

(C) The proportion of living B, NK and T cells seen after treatment (48 hrs) of fresh isolated mouse BM cells ($2\times10^6$) with BIO (1 μM), a Wnt/β-catenin signaling agonist. Cells were analyzed by flow cytometry (n=7).

(D) The proportion of living B, NK and T cells seen after treatment (48 hrs) of fresh isolated mouse BM cells ($2\times10^6$) with CJJ300 (10 μM), a TGF-β inhibitor. Cells were analyzed by flow cytometry (n=5).

(E) The proportion of living B, NK and T cells seen after treatment (48 hrs) of fresh isolated mouse BM cells ($2\times10^6$) with MSAB (4 μM), a Wnt/β-catenin signaling inhibitor. Cells were analyzed by flow cytometry (n=5).

Asterisks represent statistical significance (*p < 0.05; **p < 0.01; ***p < 0.001; ****p < 0.0001) as determined by one-way ANOVA. Error bars indicate standard error of the mean (SEM).

**Supplementary Figure 7. Fos or Klf4 target genes in B and NK cells.**

(A) Volcano plots showing differential expression of Fos or Klf4 target genes in B cells from mice immunized with G1-C versus RBD vaccine, analyzed using DESeq2. ChIP-Atlas 3.0 was used identify Fos, Klf4, and Klf6 target genes at the TSS ± 5k from M. musculus (mm10). Red dots, upregulated genes; blue, downregulated genes; gray, non-significant changes. Green dots indicate DEGs overlapping in both Fos or Klf4 target genes.

(B) Volcano plots showing differential expression of Fos or Klf4 target genes in NK cells from mice immunized with G1-C versus RBD vaccine, analyzed using DESeq2. ChIP-Atlas 3.0 was used identify Fos and Klf4 target genes at the TSS ± 5k from M. musculus (mm10). Red dots, upregulated genes; blue, downregulated genes; gray, non-significant changes. Green dots indicate DEGs overlapping in both Fos or Klf4 target genes.

**Supplementary Figure 8. G1-C sequence alignment**

(A) Sequence alignment of G1-C epitope from multiple SARS-CoV-2 variants.

**Supplementary Figure 9. Flow cytometry gating strategy for T cell responses in PBMCs and spleen**

(A) Spike-specific T cell gating strategy. Spike-specific $CD4^+$ ($CD3^+CD4^+$IFN-$\gamma^+$/IL-$2^+$/TNF$\alpha^+$) and $CD8^+$ ($CD3^+CD8^+$IFN-$\gamma^+$/IL-$2^+$/TNF$\alpha^+$) T cells were gated from the $CD3^+$ population after removing doublets and dead cells.

(B) Representative flow cytometry plots of IFN-$\gamma^+$/IL-$2^+$/TNF$\alpha^+$ $CD4^+$/$CD8^+$ T cells in PBMCs.

**Supplementary Figure 10. Flow cytometry gating strategy for B cells in spleen.**

(A) Memory B cell, germinal center B cell, and plasma cell gating strategy. Memory B ($B220^+CD21^+CD80^+$), germinal center B ($B220^+FAS^+CD38^{low}$), and plasma (Blimp-$1^+CD138^+$) cells were gated after removing doublets and dead cells. RBD-binding memory B and germinal center B cells were identified by RBD proteins. The RBD-specific B cells could recognize RBD antigen and bind to RBD[53].

(B) Representative flow cytometry plots of plasma cells, germinal center B cells and $RBD^+$ germinal center B cells.

**Supplementary Figure 11. Flow cytometry gating strategy for immune cells in bone marrow.**

(A) LT-HSC, ST-HSC, LMPP, and CMP gating strategy. LT-HSCs ($Lin^-Sca1^+c\text{-}Kit^+CD34^-CD135^-$), ST-HSCs ($Lin^-Sca1^+c\text{-}Kit^+CD34^+CD135^-$), LMPPs ($Lin^-Sca1^+c\text{-}Kit^+CD34^+CD135^+$), and CMPs ($Lin^-Sca1^-c\text{-}Kit^+CD34^+CD16/32^-$) were gated after removing doublets and dead cells.

(B) Representative flow cytometry plots of LT-HSC, ST-HSC, LMPP, and CMP.

(C) Lymphoid and myeloid lineage cell gating strategy. Lymphoid lineage cells included B ($B220^+$), NK ($CD3^-NK1.1^+$), and T ($CD3^+$) cells. Myeloid lineage cells included neutrophils ($CD45^+CD11b^+Ly6G^+$), cDCs ($CD45^+CD11b^+CD11c^+$), and monocytes ($CD45^+CD11b^+CD115^+$). All cells were gated after removing doublets and dead cells.

**Supplementary Figure 12. Flow cytometry gating strategy for CD8 T cells subsets and APCs in spleen.**

(A) Granzyme B, naïve, central memory and effector CD8 T gating strategy. Granzyme $B^+$ $CD8^+$ T cells ($CD3^+CD8^+$Granzyme $B^+$), naïve $CD8^+$ T cells ($CD3^+CD8^+CD62L^+CD44^-$), central memory $CD8^+$ T cells ($CD3^+CD8^+CD62L^+CD44^+$), and effector $CD8^+$ T cells ($CD3^+CD8^+CD62L^-CD44^+$) were gated after removing doublets and dead cells.

(B) Antigen-presenting cells gating strategy. Macrophages ($CD45^+CD11b^+F4/80^+$) and cDCs ($CD45^+CD11b^+CD11c^+$) were gated after removing doublets and dead cells.
